# Physiologic RNA Targets and Refined Sequence Specificity of Coronavirus EndoU

**DOI:** 10.1101/2020.05.20.064436

**Authors:** Rachel Ancar, Yize Li, Eveline Kindler, Daphne A. Cooper, Monica Ransom, Volker Thiel, Susan R. Weiss, Jay R. Hesselberth, David J. Barton

## Abstract

Coronavirus EndoU inhibits dsRNA-activated antiviral responses; however, the physiologic RNA substrates of EndoU are unknown. In this study, we used mouse hepatitis virus (MHV)-infected bone-marrow-derived macrophage (BMM) and cyclic phosphate cDNA sequencing to identify the RNA targets of EndoU. EndoU targeted viral RNA, cleaving the 3′ side of pyrimidines with a strong preference for U^⬇^A and C^⬇^A sequences (endoY^⬇^A). EndoU-dependent cleavage was detected in every region of MHV RNA, from the 5′ NTR to the 3′ NTR, including transcriptional regulatory sequences (TRS). Cleavage at two CA dinucleotides immediately adjacent to the MHV poly(A) tail suggest a mechanism to suppress negative-strand RNA synthesis and the accumulation of viral dsRNA. MHV with EndoU (EndoU^mut^) or 2′-5′ phosphodiesterase (PDE^mut^) mutations provoked the activation of RNase L in BMM, with corresponding cleavage of RNAs by RNase L. The physiologic targets of EndoU are viral RNA templates required for negative-strand RNA synthesis and dsRNA accumulation.

**Impact:** Coronavirus EndoU cleaves U^⬇^A and C^⬇^A sequences (endoY^⬇^A) within viral (+) strand RNA to evade dsRNA-activated host responses.

## INTRODUCTION

Viruses in the order *Nidovirales* express a virus-encoded endoribonuclease, NendoU (1). NendoU is unique to nidoviruses (2), including viruses of the *Coronaviridae* and *Arteriviridae* families. Nidoviruses that express NendoU have vertebrate hosts whereas nidoviruses of crustaceans (*Roniviridae*), and RNA viruses outside the *Nidovirales* order, do not encode this protein. The precise role(s) of NendoU in virus replication remain enigmatic; however, significant progress has been made in recent years to elucidate the contributions of NendoU to virus replication and pathogenesis. The SARS-CoV-2 pandemic underscores the importance of understanding host-pathogen interactions, including the immunomodulatory functions of EndoU (3).

Arterivirus (nsp11) and coronavirus (nsp15) EndoU proteins have been characterized by genomic (2), structural (4–6) and biochemical studies (6–8). EndoU is encoded near the 3′ end of ORF1b (Fig. 1A, schematic of MHV genome) (2). Mouse hepatitis virus (MHV), a well-studied coronavirus, has a single-stranded positive-sense RNA genome 31.1 kb in length. MHV RNA, like other coronaviruses, is 5′ capped and 3′ polyadenylated. Upon infection, the ORF1a and ORF1b regions of MHV RNA are translated into two polyproteins (ORF1a and ORF1ab) through a frame shifting mechanism (9). MHV proteins nsp1-nsp16 are produced via proteolytic processing of the ORF1a and ORF1ab polyproteins. EndoU is the nsp15 protein of MHV (Fig. 1A, schematic of MHV RNA genome). Other proteins from the ORF1a/1b region of the RNA genome include viral proteases and components of the viral replicase (nsp12 is the RdRP, nsp13 is a helicase, nsp14 is a 3′⃗5′ exonuclease and a N7-methyl transferase, and nsp16 is a 2’-O-methyl transferase). An H277A mutation in nsp15 disables the catalytic activity of EndoU (Fig. 1A, EndoU^mut^).

**Figure 1.**
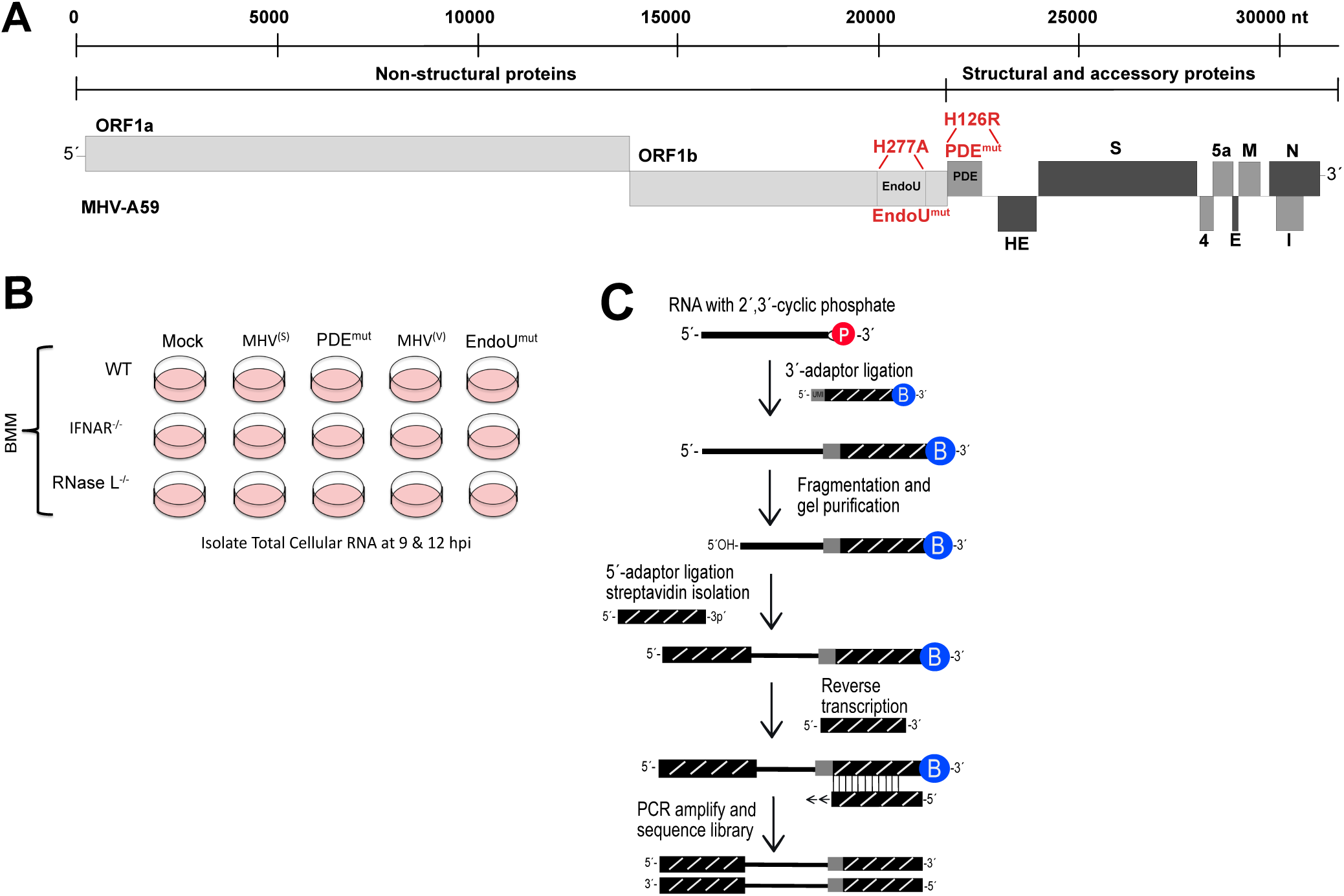
Coronavirus RNA genome and Experimental Approach. (A) MHV RNA genome highlighting two mutations: His to Arg mutation in the MHV phosphodiesterase domain active site (PDE^H126R^), and a His to Ala mutation in the MHV EndoU-domain active site (EndoU^H277A^) (26, 38). MHV proteins are categorized as nonstructural (light grey), accessory (dark gray) and structural (black). (B) Bone marrow-derived macrophage (BMM) from WT, IFNAR^-/-^, and RNase L^-/-^ mice were mock-infected or infected with WT MHV [MHV^(S)^ and MHV^(V)^], the PDE^mut^, or EndoU^mut^ for 9 and 12 hours (26, 29), after which total cellular RNA was isolated for cyclic phosphate sequencing. (C) Schematic of cyclic phosphate sequencing, protocol adapted from Schutz et al., 2011 (34).

Coronavirus RNA replication and RNA transcription are mediated by the replicase expressed from the ORF1a/1b region of the genome, with assistance of the nucleocapsid protein (10). Both RNA replication and RNA transcription occur within membrane-anchored replication organelles in the cytoplasm of infected cells (11–13). MHV RNA replication involves negative-strand RNA synthesis, wherein the positive-strand viral RNA genome is copied into a genome-length negative-strand RNA intermediate, which is subsequently used as a template to make new positive-strand RNA genomes. MHV RNA transcription involves the synthesis of subgenomic (sg) negative-strand RNAs from the viral RNA genome via discontinuous transcription mechanisms and subsequent synthesis of sg mRNAs (14, 15). Intergenic transcriptional regulatory sequences (TRS) within MHV RNA guide discontinuous transcription mechanisms (16), leading to the production of sg negative-strand RNAs, which function as templates for the synthesis of sg mRNAs. A nested set of 3′ co-terminal sg mRNAs (sg mRNA2 to sg mRNA7) is used to express each of the remaining viral proteins [phosphodiesterase (PDE) from sg mRNA2a; spike (S) from sg mRNA3, and so forth). Hemagglutinin-esterase (HE) is an unexpressed pseudogene in MHV A59 due to a TRS mutation that prevents the expression of mRNA2b, as well as a nonsense mutation at codon 15 (17–20). EndoU co-localizes with viral RNA replication and RNA transcription machinery at membrane-anchored replication organelles (21, 22). Co-localization of EndoU with viral RNA synthesis machinery may influence the RNAs targeted by EndoU. Furthermore, coronavirus nsp16, a 2′-O-ribose-methyltransferase (2′-O’MT), could potentially modify RNA substrates to make them resistant to cleavage by EndoU (1). These studies suggest that viral RNA stability may be regulated by nsp15 (EndoU) and nsp16 (2′-O’MT).

Intriguingly, neither EndoU (nsp15) nor 2′-O’MT (nsp16) enzyme activities are required for virus replication in transformed cells in culture (23–25); rather, these enzymes counteract dsRNA-activated antiviral responses (22, 25, 26). EndoU catalytic activity prevents the activation of dsRNA-dependent antiviral innate immune pathways (22, 26), including Type I and Type III IFN responses, PKR and OAS-RNase L (27). EndoU-deficient viruses can replicate in IFNAR^-/-^ cells or cells lacking PKR and RNase L (PKR^-/-^ & RNase L^-/-^) (22, 26, 27). In addition to EndoU, coronavirus NS2, a 2′-5′ PDE, prevents activation of RNase L (28–30). Thus, there are two pathways by which MHV prevents activation of OAS-RNase L suggesting this pathway is crucial for antiviral defense. While coronavirus EndoU inhibits dsRNA-activated antiviral responses within virus-infected cells, it is unclear how it achieves this because the physiologically relevant targets of EndoU have not been defined.

In this study, we used MHV-infected bone marrow-derived macrophage (BMM) and cyclic phosphate cDNA sequencing to identify the host and viral RNA targets of EndoU. Cyclic phosphate cDNA sequencing reveals the location and frequency of endoribonuclease cleavage sites within host and viral RNAs (31–34). We exploited wildtype and mutant forms of MHV (wt MHV, PDE^mut^, and EndoU^mut^) along with wildtype and mutant forms of BMM (wt BMM, IFNAR^-/-^ and RNase L^-/-^) to distinguish between EndoU-dependent cleavage sites and RNase L-dependent cleavage sites within host and viral RNAs.

## MATERIALS AND METHODS

### Viruses

Wildtype Mouse Hepatitis Virus A59 from Volker Thiel [MHV^(V)^] (35–37) and Susan Weiss [MHV^(S)^] (38) were used, along with a mutant derivative of each. An H277A mutation in nsp15 rendered an EndoU-deficient mutant (EndoU^mut^) from MHV^(V)^ (26). An H126R mutation in NS2 rendered a phosphodiesterase mutant (PDE^mut^) from MHV^(S)^ (38).

### Murine bone marrow-derived macrophages

Bone marrow-derived macrophage (BMM) from WT, IFNAR^-/-^ and RNase L^-/-^ C57BL/6 mice were obtained as previously described (26). Progenitor cells were isolated from the hind limbs of 8-12 week old mice, passed through a cell strainer and RBCs were lysed using 1 ml of lysis buffer (0.15 M NH_4_Cl, 1 mM KHCO_3_, 0.1 mM EDTA). Cells were washed 3x with PBS and cultured in macrophage medium (Iscove’s Modified Dulbecco’s Medium, 5-10% M-CSF (L929-supernatant), 0.1% 50 mM 2-mercaptoethanol). Adherent BMM were harvested at 7 dpi.

### Virus infection

BMM were infected with MHV^(S)^, MHV^(V)^, EndoU^mut^, and PDE^mut^ at an MOI of 1 PFU per cell at 37°C as previously described (26). At 9 and 12 hours post-infection (hpi), supernatant was harvested for virus titration and cells were lysed in Trizol (Invitrogen). MHV in the supernatant was quantified by standard plaque assay on L2 cells.

### Cyclic phosphate cDNA sequencing

Total RNA was extracted from cell lysates and split equally for cyclic phosphate and RNAseq library preparations. Cyclic phosphate cDNA libraries were prepared by DNase treating the total RNA for 30 min followed by ethanol precipitation with 20 μg of glycogen and ligation with 50 μM 3’-linker in 30 μl final volume. The ligation reactions were conducted using 15 pmol of RtcB ligase (NEB), 1x RtcB buffer (NEB), 100 μM GTP, 1 mM MnCl_2_, 20 units of RNase inhibitor (Enzymatics) at 37°C for 2 h. Samples were ethanol precipitated with 20 μg of glycogen and resuspended in 10 μl of RNase free H_2_O for chemical fragmentation (Ambion Fragmentation Reagent) at 65°C for 4 min. Samples were then denatured in 1 volume of stop dye (95% formamide, 0.01% xylene cyanol / bromophenol blue), heated to 65°C for 5 min and separated on a 6 % polyacrylamide TBE–urea gel. Gels were stained with SYBR Gold (Invitrogen) and visualized to excise RNA larger than adapter (∼100-1000 bp). RNA was eluted from the gel slices with 2 h incubation at 40°C in 0.3 M sodium acetate, pH 5.2, 1 mM EDTA, pH 8.0 followed by gentle mixing overnight at 4°C. Eluted RNA was recovered by ethanol precipitation with 20 μg of glycogen and resuspended in 12 μl of RNase free H_2_O. RNA was ligated to 50 μM 5’-linker in 20 μl final volume. The ligation reactions were conducted using 15 pmol of RtcB (NEB), 1x RtcB buffer (NEB), 100 μM GTP, 1 mM MnCl_2_, 20 units of RNase inhibitor (Enzymatics) at 37°C for 2 h followed by ethanol precipitation with 20 μg of glycogen and resuspended in 100 μl of RNase free H_2_O. Ligated RNAs were purified using 25 μl of magnetic Streptavidin beads (Invitrogen) washed three times with 100 μl of B&W buffer [5 mM Tris-HCl (pH 7.5), 0.5 mM EDTA, 1 M NaCl] supplemented with 0.1% Tween 20, twice with 100 μl of solution A (0.1 M RNase free NaOH, 0.05 M RNase free NaCl), and twice with 100 μl of solution B (0.1 M RNase free NaCl). Washed beads were resuspended in 2x B&W buffer [10 mM Tris-HCl (pH 7.5), 1 mM EDTA, 2 M NaCl], with 20 units of RNase inhibitor (Enzymatics) and the RNA solution was added to the beads and incubated with rotation for 15 min at room temperature. After incubation, the beads were washed three times with 100 μl of 1x B&W buffer before resuspending the beads in 20 μl of 25 mM biotin in elution buffer (Omega BioTek). The beads were incubated at room temperature for 15 min with occasional mixing. After binding the beads to the magnet, the supernatant was collected. The elution process was repeated once for a final volume for 40 μl of eluted RNA. cDNA was prepared using 5 μM of an Illumina-compatible primer complimentary to the 3’-linker, 20 ul of eluted RNA, and Protoscript II RT (NEB). 10 μl of cDNA was PCR amplified for 18 cycles with Illumina TruSeq primers and Phusion DNA polymerase. PCR reactions were purified with AMPure XP beads (Beckman Coulter). Indexed libraries were quantified by Qubit (Invitrogen). Library quality was assessed on a 4200 TapeStation System Instrument (Agilent Technologies) using a D100 ScreenTape assay, mixed to a final concentration of 1–10 nM and sequenced on an Illumina HiSeq in a 50 cycle run.

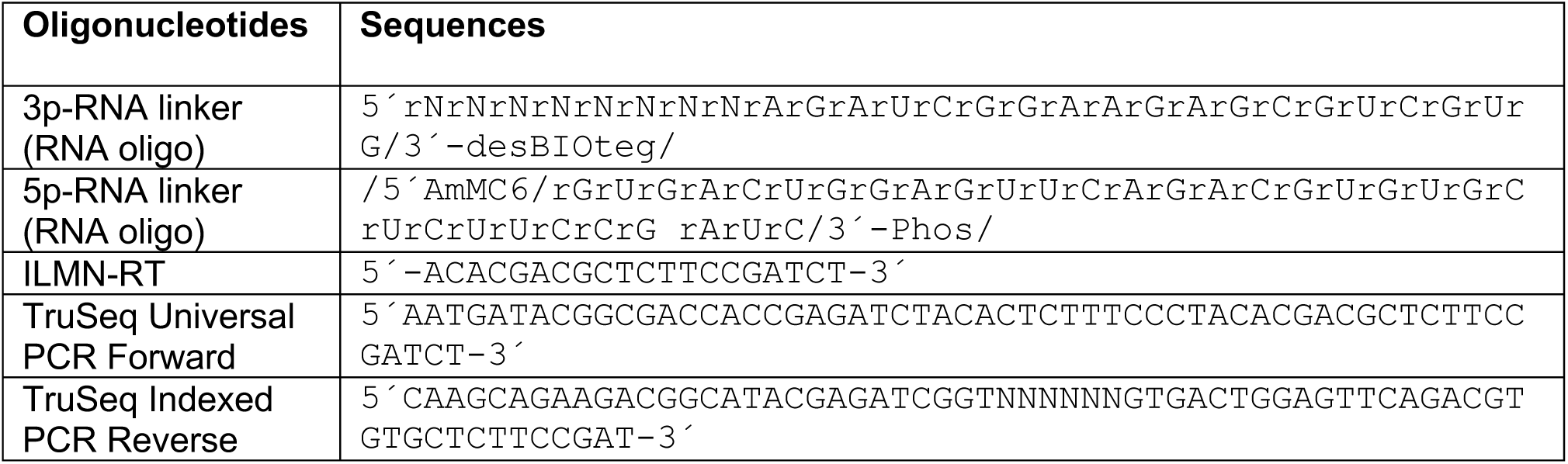

### Stranded RNAseq

Total RNA was enriched for polyadenylated mRNA using oligo-dT magnetic beads (Ambion). cDNA was generated from the enriched polyA^+^ mRNAs after fragmentation in 2.2x SuperScript IV reverse transcriptase buffer at 94°C for 3 min. After immediately cooling on ice, RT reaction with SuperScript IV RT (Thermo Fisher Scientific) was performed per manufacture’s recommendations with 150 ng of random primers (Thermo Fisher Scientific) in 20 μl final volume. cDNA:RNA hybrids were purified using MyOne Silane beads (Thermo Fischer Scientific) per manufacture’s recommendations and eluted in 18 μl of RNase free H_2_O. Second-strand cDNA was then generated using RNase H and *E. coli* DNA Polymerase (Enzymatics) with dUTP incorporation (1x NEB buffer 2, 100 μM dATP, dCTP, dGTP, 200 μM dUTP, 2.5 units of RNase H, 30 units of DNA polymerase) in 100 μl final volume at 15°C for 2.5 hrs. cDNA was purified with Silane beads and eluted in 52 μl of RNase free H_2_O as input for end repair reaction using End Repair Module (NEB) following manufacturer’s recommendations. A-tailing reaction (50 μl final volume) performed with Klenow fragment (minus 3’-5’exonuclease activity, Enzymatics) and end repaired Silane purified cDNA eluted in in 32 μl of RNase free H_2_O (1x NEB buffer 2, 200 uM dATP, 15 units Klenow fragment) at 37°C for 30 min. Reaction products were purified with 1.8x AMPure XP beads (Beckman Coulter) and eluted in 10 μl of RNase free H_2_O. Purified cDNA was ligated to 40 nM of annealed Illumina TruSeq Universal adaptors in 50μl final volume reaction for 30 min at 25°C (40 nM adaptors, 1X Rapid Ligation Buffer (Enzymatics), 3000 units of T4 DNA ligase (Enzymatics). Reaction products were purified with AMPure XP beads and eluted in 12 μl of RNase free H_2_O. USER enzyme (NEB) was used to degrade the dUTP-containing strand by adding 1 unit of USER to purified cDNA and incubating for 30 min at 37°C. Reactions were used directly in PCR amplification with Illumina TruSeq primers and Phusion DNA polymerase with 10 μl of input for 18 cycles. Libraries were size selected from 200 – 700 bp using AMPure XP beads, quantified by Qubit (Invitrogen), and mixed to a final concentration of 4 nM. Library quality was assessed on a 4200 TapeStation System Instrument (Agilent Technologies) using a D100 ScreenTape assay and sequenced on an Illumina NovaSEQ 6000 in a paired end 150 cycle run.

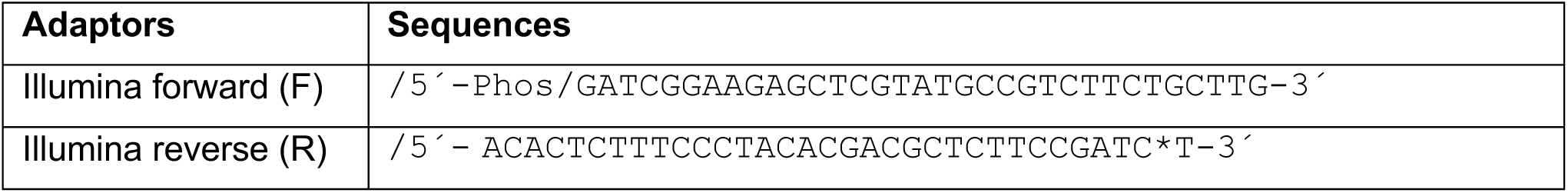

### Computational analyses of next generation sequencing data

#### Processing and analysis of cyclic phosphate cDNA libraries

Unique molecular identifier (UMI) sequences were extracted and added to FASTQ reads using UMI-tools (v0.5.4) (39). Only read 1 was used from the second experiment, to adhere with the analysis process applied for experiment 1. FASTQ reads were then aligned to the MHV genome alone (GenBank accession: NC_001846.1) and a combined reference including the MHV genome, *Mus musculus* rRNA and U6 snRNA references (GenBank accession numbers: NR_003278.3, NR_003279.1, NR_003280.2, NC_000074.6, NR_003027.2), and annotated ORFs from the Mouse ORFeome collection (MGC full-cds collection for Mus musculus) using Bowtie version 2 (v2.3.2) (40). Aligned reads were de-duplicated using UMI-tools to remove PCR duplicated reads. De-duplicated reads were converted to bedGraph format using BEDTools (v2.26.0) to report the number of reads at each single base cleavage position, including for sense and antisense sequences for the MHV aligned reads (41). Reads at each cleavage position were normalized by library size.

To identify signal dependent on the presence of a specific endoribonuclease, normalized counts at each cleavage position in RNase L^-/-^ or EndoU^mut^ libraries were subtracted from the signal in libraries with wild type RNase L or EndoU activity, to remove signal that occurred in the absence of either endoribonuclease. The difference in cleavage activity at each position in the absence of RNase L or EndoU was determined by calculating the log2 fold change. The frequency of cleavage at particular dinucleotides was determined by quantifying the sum of reads assigned to each of the 16 possible dinucleotides divided by total number of aligned reads in the library. Dinucleotide enrichment was determined by calculating the frequency of cleavage at each dinucleotide in the MHV genomic sequence and determining the log2 fold enrichment of the observed (experimental) frequencies compared to the expected (background) frequencies. Significance of enrichment was calculated using the Fisher’s exact test to compare the odds ratio of obtaining a specific dinucleotide in the expected data to the observed data.

#### RNAseq alignment, annotation, and differential expression analysis

Illumina adaptor sequences were trimmed from FASTQ reads using Cutadapt (v 1.16) and sequences shorter than 20 nucleotides were discarded (42). Trimmed reads were aligned to a combined MHV (GenBank accession: NC_001846.1) and *Mus musculus* genome reference (Ensemble GRCm38.p6) and bedGraph coverage files were generated from each alignment using STAR (v 2.7.1a) (43). Read fragments were assigned and counted using featureCounts (from subread v 1.6.2) and a combined MHV and *Mus musculus* GTF (Ensemble GRCm38.p6) file for gene annotation (44). The MHV GTF file included the genomic positions of the combined ORF1a/b non-structural proteins and each of the structural and accessory proteins. Gene counts were normalized using DESeq2 media of ratios method to account for sequencing depth and RNA composition (45). For downstream differential expression analysis, trimmed reads were also aligned to the *Mus musculus* complete transcriptome reference (Gencode GRCm38.p6) using Salmon (v.0.14.1) (46). Transcript abundance files were used for differential expression analysis with DESeq2 after importing with tximport and counts normalized by the media of ratios method were used for data visualization (47). Genes with an FDR < 0.05 were called significant and used to generate volcano plots with the EnhancedVolcano package and z-transformed counts were used to generate heatmaps with the ComplexHeatmap package (48, 49). For gene functional category enrichment analyses, topGO was used to determine significant enrichment (weightFish/p > 0.01) by using non-differentially expressed genes (< 2 or < −2 log2 fold change and FDR < 0.01) as the background to determine the categories enriched in differentially expressed genes. topGO employs conditional enrichment analysis, which takes the nested structure of GO terms into account to reduce redundancy in enrichment results (50).

#### Motif analysis

To visualize the cleavage sequence preferences for RNase L and EndoU, the top 1% of either RNase L-or EndoU-dependent sites from subtractive analysis, as described above, were selected. Using BEDTools, 3 bps were added upstream and downstream of the selected positions and a FASTA file was generated from the 6-base pair sequences. Meme was used to determine the sequence preference enrichment and graphed using ggseqlogo (51, 52).

#### UA scoring

UA sequences in the MHV genome were designated as predominantly cleaved 3’ of U (consistent with EndoU targeting) or A (consistent with RNase L targeting). All UA dinucleotides in the MHV genome with > 30 cyclic phosphate counts at either position of cleavage in the dinucleotide were selected and the ratio of normalized counts in each position was calculated (RNase L / EndoU). If the ratio was > 1, the position was scored as a UA^⬇^ site and if the ratio was < 1 the position was scored as a U^⬇^A.

#### Regional MHV cleavage analysis and abundance normalization

The normalized counts in each MHV genomic region (all the genes and ORFs shown in Figure 1A, in addition to the 5’and 3’-UTR and body TRS regions) were summed to calculate the total cyclic phosphate reads per region. Size correction was performed by dividing the sum of cyclic phosphate counts in each region by the length of the region in bases. To normalize the cyclic phosphate data by RNA abundance, stranded bedGraph files were generated from the bam files produced by STAR alignment of the RNAseq libraries. At each position with both cyclic phosphate and RNAseq data in the MHV genome, the cyclic phosphate counts were divided by the normalized (reads per million mapped reads) RNAseq counts to generate an abundance normalized cyclic phosphate value.

#### TRS analysis

In the above analyses, RNAseq reads mapping to the viral genome were not distinguished by alignment to genomic RNA or subgenomic mRNAs. To assign RNAseq reads to subgenomic mRNAs, we employed an analysis similar to that described in Irigoyen et al., 2016 (53) to identify leader/body chimeric reads (subgenomic mRNAs). We parsed the bam files generated from STAR alignment of the RNAseq libraries to the combined mouse and MHV genome for reads containing the 11 nucleotides of the leader sequence, UUUAAAUCUAA (GenBank accession: NC_001846.1, nt 54 – 65) before the leader TRS sequence. The positions in the reference where the read alignment starts and ends after the leader sequence were extracted to obtain an interval of alignment for the sequence downstream of the leader/body transition. The intervals for each chimeric read were intersected with the intervals of each canonical subgenomic mRNA using valr (54), with the requirement of at least 30 nt of overlap, to assign each chimeric read to an mRNA: 65 to 21746 (mRNA 1), 21747 to 23921 (mRNA 2), 23922 to 27934 (mRNA 3), 27935 to 28317 (mRNA 4), 28318 to 28959 (mRNA 5), 28958 to 29654 (mRNA 6), 29655 to 31334 (mRNA 7) (54). The number of reads assigned to each mRNA were counted and normalized to either the total sum of mRNAs per library or reads per million.

#### SNP analysis

Variant calling analysis was performed using bcftools (v1.9) to generate genotype likelihoods from the RNAseq bam files for MHV aligned reads, followed by SNP calling/indel calling to generate VCF files (55). The generated VCF files were filtered using bcftools with parameters -s LOWQUAL -e %QUAL<30 || DP>20’ to identify low quality sites with less than 20 quality score or 30 base pairs of read depth.

### Data Deposition

Raw and processed sequencing data are available at NCBI GEO: GSE147852.

### Bioinformatics Pipeline

Code for all described analyses are available at https://github.com/hesselberthlab/endoU in the form of scripts, a data processing pipeline, and analysis package.

## RESULTS

Products of cleavage by coronavirus EndoU have 2′,3′-cyclic phosphate termini (1), effectively marking the location of cleavage within host and viral RNAs. Thus, in this study, we used cyclic phosphate cDNA sequencing to monitor the frequency and location of endoribonuclease cleavage sites in RNA from MHV-infected bone marrow-derived macrophage (BMM) (Fig. 1). Wildtype and mutant MHVs [wt MHV^(V)^, wt MHV^(S)^, PDE^mut^, and EndoU^mut^] along with BMM derived from wildtype and particular knockout C57BL/6 mice [WT, IFNAR^-/-^ and RNase L^-/-^ BMM] were used to distinguish between EndoU-dependent cleavage sites and RNase L-dependent cleavage sites (Fig. 1B). A pair of wt and mutant viruses derived from each isolate were used (Figs. 1A and 1B): wt MHV from Susan Weiss’ lab designated MHV^(S)^ and a phosphodiesterase mutant designated PDE^mut^ (28–30, 38); wt MHV from Volker Thiel’s lab designated MHV^(V)^ and an EndoU mutant designated EndoU^mut^ (26). Total cellular RNA was isolated from cells at 9 and 12 hpi, times when coronavirus NS2 PDE and nsp15 EndoU activities prevent dsRNA-dependent antiviral responses (26), including the OAS/RNase L pathway (28–30). Under these experimental conditions (Fig. 1B), we expect that RNase L activity will be increased within PDE^mut^-infected and EndoU^mut^-infected WT BMM, as compared to MHV^(S)^-infected and MHV^(V)^-infected WT BMM. Furthermore, we expect that EndoU activity will be evident within MHV^(S)^-infected and MHV^(V)^-infected BMM, as compared to EndoU^mut^-infected BMM.

Cyclic phosphate cDNA libraries were prepared using total cellular RNA from 9 and 12 hpi (Fig. 1C). The RNA ligase RtcB was used to ligate a 3′ adaptor to RNA fragments containing a cyclic phosphate. The 3′ adaptor has a biotin moiety and a unique molecular identifier to enumerate cleavage sites (56). A 5′ adaptor was ligated to the RNA samples, followed by reverse transcription, PCR amplification and Illumina sequencing. Analysis of DNA sequences revealed the frequency and location of endoribonuclease cleavage sites in host and viral RNAs. Figures 2-8 in the body of this manuscript correspond to data from the experiment outlined here (Fig. 1). Replicate data from infections by wt and mutant MHV [wt MHV^(S)^, PDE^mut^, and EndoU^mut^] in wt and RNase L^-/-^ BMM yield similar outcomes (Figures S8 and S9).

**Figure 2.**
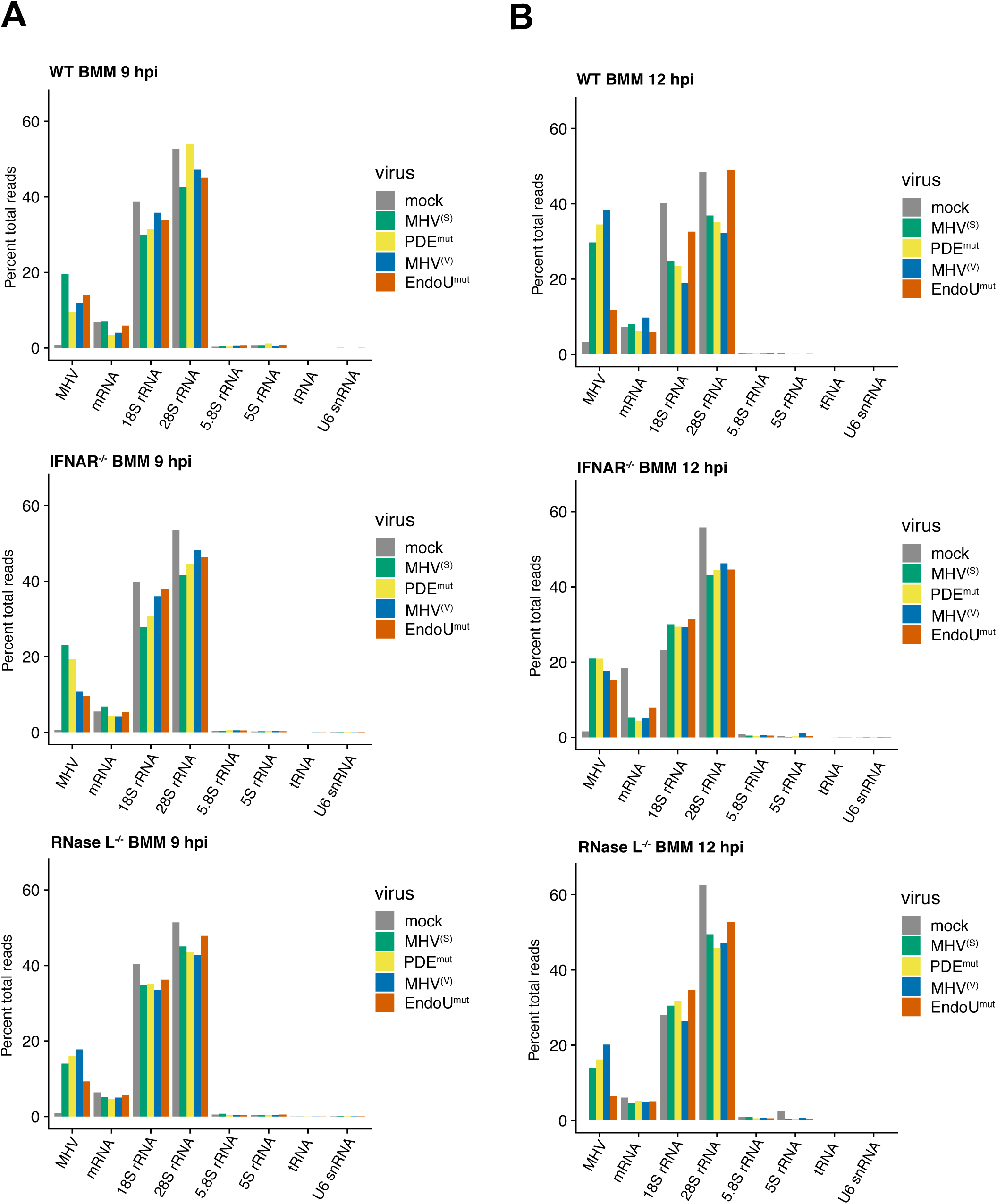
Frequency of endoribonuclease cleavage in host and viral RNAs. (A and B) Normalized cyclic phosphate cDNA reads ([reads at each position / total reads in library]) mapped to host and viral RNAs at 9 and 12 hpi in WT, IFNAR^-/-^, and RNase L^-/-^ bone marrow macrophages (BMM).

### Endoribonuclease cleavage sites in host and viral RNAs

Endoribonuclease cleavage sites were detected in both host and viral RNAs (Fig. 2). The frequency of cleavage sites in individual RNAs was normalized to percent total cDNA reads in each library, allowing for quantitative comparisons between individual RNAs in each sample and between RNAs across distinct samples. The vast majority of cleavage sites were detected in MHV RNA, cellular mRNA and ribosomal RNAs (18S, 28S 5.8S and 5S rRNAs), with a smaller portion of cleavage sites in tRNAs and U6 snRNA (Fig. 2). We can attribute cleavage sites in cellular RNAs to specific endoribonucleases in some cases, but not others. For instance, U6 snRNA had 3′-terminal cyclic phosphates (Fig. S1A) attributed to the nucleolytic activity of C16orf57/USB1 (32, 57, 58). Ribosomal RNAs accounted for ∼50-80% of the cleavage sites detected in each library (Fig. 2). The majority of cleavage sites within rRNAs are the result of unspecified endoribonucleases, along with some RNase L-dependent cleavage sites (31, 32), as described below. Cellular mRNAs accounted for ∼5% of endoribonuclease cleavage sites in each cDNA library (Fig. 2); however, the numbers of cleavage sites within individual cellular mRNAs were too low to definitively attribute to one or another endoribonuclease.

Cleavage sites in MHV RNA were found predominantly in the positive-strand of viral RNA, ranging from 10% to 40% of all cleavage sites in each library (Fig. 2). Very few reads were detected in the MHV negative-strand RNA (Fig. S1B, S11C-D). As described below, we attribute cleavage sites in MHV RNA to specific endoribonucleases, including EndoU and RNase L.

In WT BMM, we captured more reads per library mapping to MHV RNA at 12 hpi as compared to 9 hpi, except in cells infected with EndoU^mut^ MHV. However, in IFNAR^-/-^ and RNase L^-/-^ BMM, capture of MHV RNA was similar between 9 and 12 hpi (Fig 2). Across all cell types the relative amount of host RNAs captured at 9 and 12 hpi were similar and in agreement with capture frequencies from uninfected and virus-infected cells previously reported (25, 26, 46). Data from an independent experiment revealed similar outcomes, with 10% to 30% of all cleavage sites in MHV RNA, 5% to 10% of cleavage sites in cellular mRNA and more than 60% of cleavage sites in ribosomal RNAs (Fig. S8A).

### Frequency, location and sequence specificity of cleavage sites in MHV RNA

Metal-ion-independent endoribonucleases have characteristic specificities: e.g. RNase A family members (RNase 1-8) cleave RNA 3′ of pyrimidines while RNase L cleaves RNA 3′ of UpN^⬇^ dinucleotides (UA^⬇^, UU^⬇^ > UG^⬇^) (32, 59). EndoU is reported to cleave RNA 3′ of pyrimidines in vitro (1, 8, 60); however, physiologically relevant targets of EndoU have not been defined.

We detected endoribonuclease cleavage sites throughout MHV RNA, under all experimental conditions (Fig. 3). The frequency of cleavage at each base of MHV RNA ranged from ∼ 0.00 to 0.2% of all cDNA reads in each library (Fig. 3, y-axis). Peaks of cleavage approaching 0.2% of all cDNA reads in each library (corresponding to 1 in 500 cleavage sites across all RNAs in each cDNA library) are present at particular sites in the N gene open reading frame, near the 3′ terminus of MHV RNA (Fig. 3, WT BMM, PDE^mut^ and EndoU^mut^). Typically, when measurable cleavage was detected at a particular base in MHV RNA at 9 hpi, measurable cleavage was also detected at that same site at 12 hpi, often with increased abundance (Fig. 3, overlapping orange and blue lines at each base for 9 and 12 hpi).

**Figure 3.**
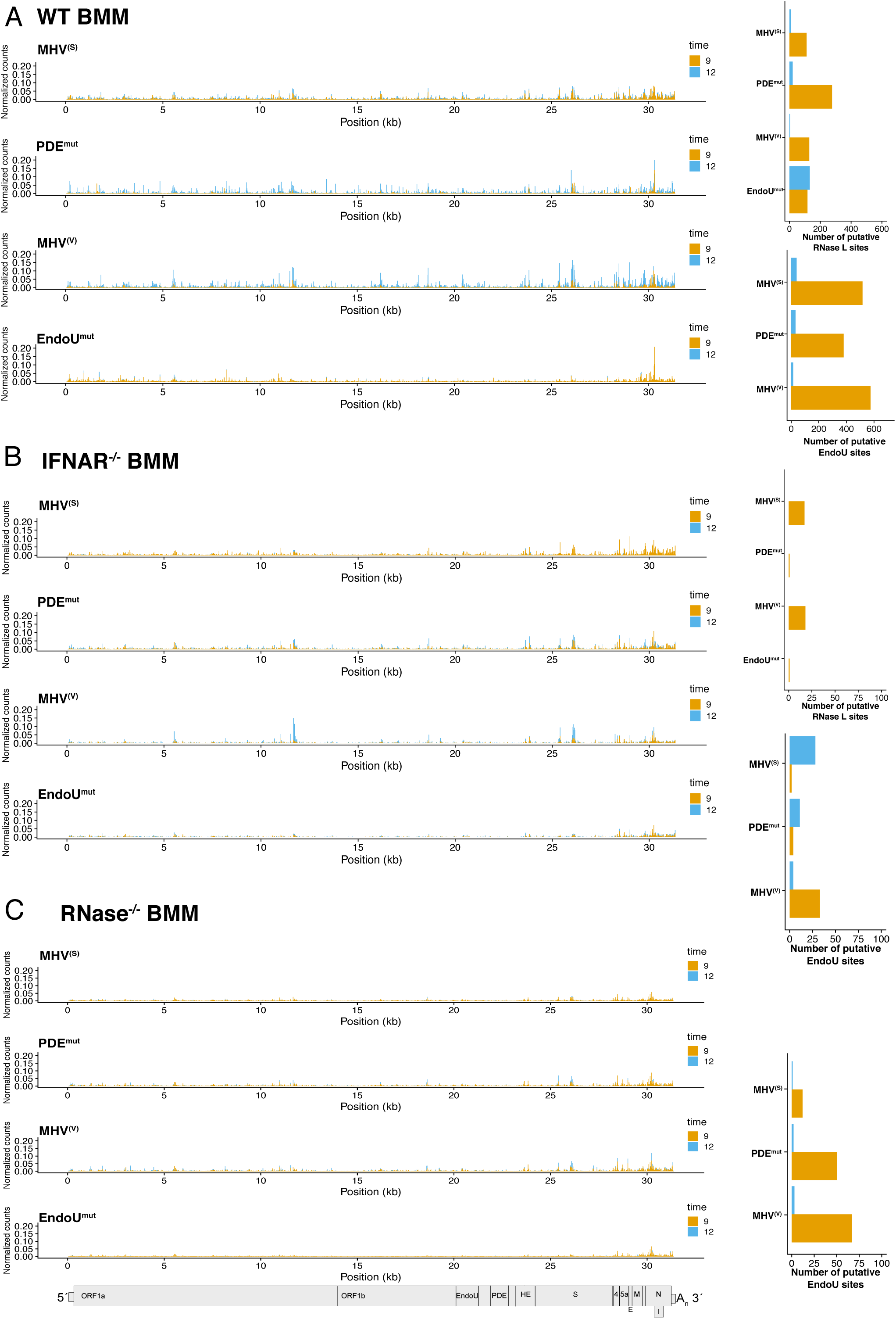
Frequency and location of endoribonuclease cleavage sites in MHV genomic RNA. (A and B) Normalized cyclic phosphate cDNA reads captured at each position along the MHV genomic RNA at 9 and 12 hpi with MHV^(S),^ MHV^(V)^, PDE^mut^, and EndoU^mut^ virus in (A) WT BMM, (B) IFNAR^-/-^, and (C) RNase L^-/-^ BMM. Putative cleavage sites attributed to EndoU or RNase L were calculated from RNase L- or EndoU-dependent signal generated by subtracting signal from each captured position that occurs in the absence of either enzyme (RNase L^-/-^ BMM or during EndoU^mut^ infection). These data were then filtered for sites with reads representing at least 0.01 % of total reads in the library. At each of these positions, the log_2_ fold change in signal when either RNase L or EndoU were absent was calculated and sites with ≥ 2.5 fold change were designated putative RNase L or EndoU sites.

The sequence specificity of cleavage sites in MHV RNA revealed profound differences in the endoribonuclease activities present within WT BMM cells infected with WT and mutant viruses (Fig. 4). Distinct RNase L-dependent and EndoU-dependent cleavage specificities were evident (Fig. 4). The sequence specificity of endoribonuclease cleavage sites was assessed in two registers: positions −2 to −1 of cleavage (Fig. 4A, B and C) and positions −1 to +1 of cleavage (Fig. 4D, E and F). WT MHV RNA was cleaved 3′ of pyrimidines in WT BMM [Fig. 4A, MHV^(S)^ and MHV^(V)^], with a notable preference for cleavage between U^⬇^A and C^⬇^A sequences (Fig. 4D, E and F). This pattern of pyrimidine specific cleavage between U^⬇^A and C^⬇^A sequences was lost in Endo^mut^-infected WT BMM (Fig. 4A and 4D). Similar patterns of cleavage were evident in an independent experiment (Fig. S8C-D).

**Figure 4.**
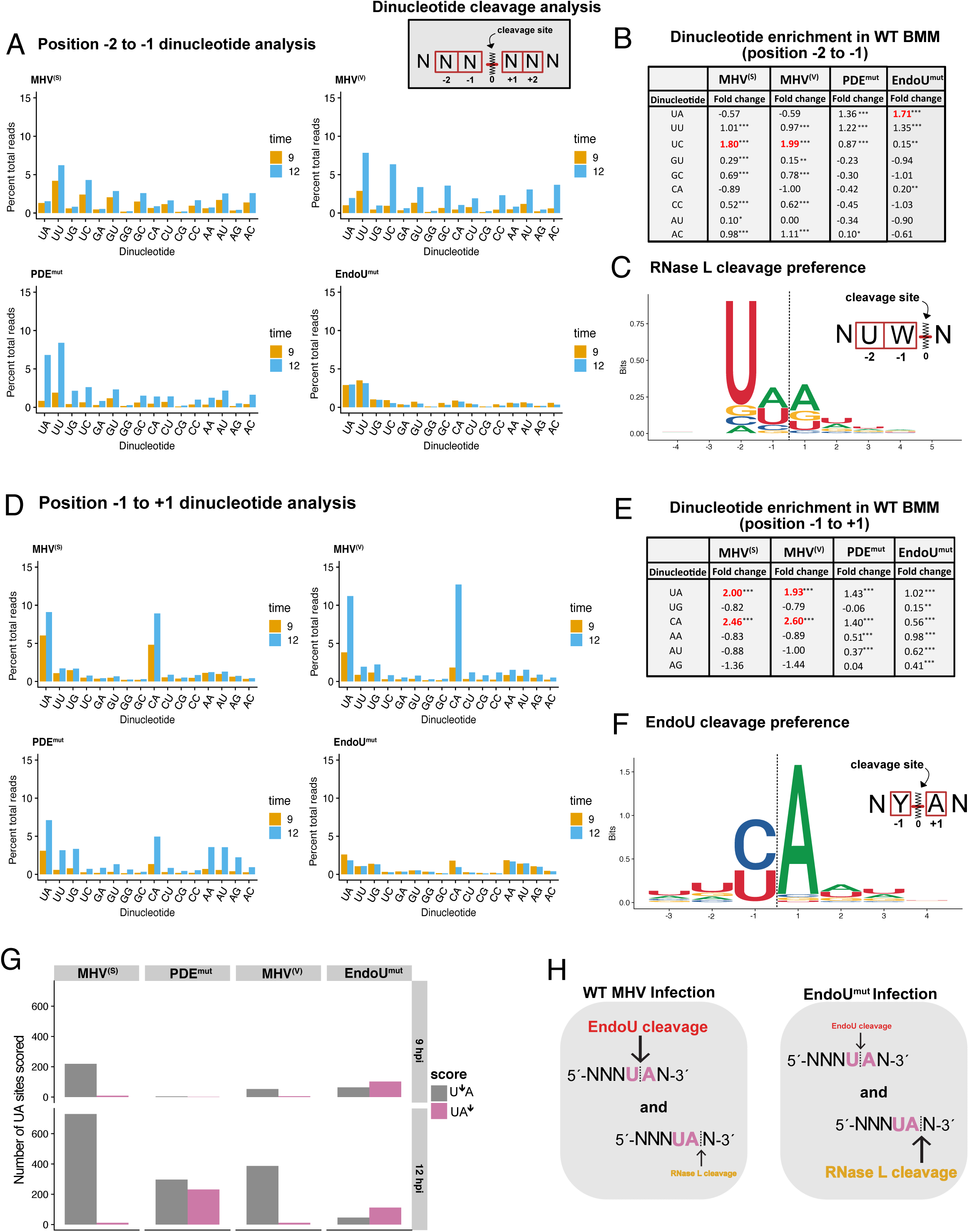
Dinucleotide endoribonuclease cleavage preference of MHV genomic RNA. (A and D) Dinucleotide specificity analysis for cleavage in MHV genomic RNA by percent total cDNA reads captured at each 3’-dinucleotide in WT BMM at 9 and 12 hpi for (A) Dinucleotide analysis for positions −2: −1 and (D) Dinucleotide analysis for positions −1:+1 from captured cleavage position (0 position). (B and E). Dinucleotide enrichment for dinucleotide positions from −2: −1 (B) or −1: +1 (E) for each condition of viral infection at 12 hpi in WT BMM by comparing the frequency of dinucleotide capture in experimental conditions to the frequency of occurrence for each dinucleotide in the MHV genomic RNA sequence (control). Significant enrichment was determined by adjusted p-value (q) for fold change ([log2(experiment / control)]). (<0.02*, <0.0001**, <1×10^8***^). Only dinucleotides with positive enrichment are shown. (C and F) Sequence logos for the 6 bases surrounding the cleavage site for position −2:-1 (C) or −1:+1 (F). Logos generated from the top 1% of either RNase L (215 sites) or EndoU-dependent cleavages (306 sites). (G) UA cleavage scoring analysis. All UA sequences in the MHV genomic RNA with ≥ 30 cyclic phosphate counts in either the UA^⬇^ or U^⬇^A cleavage position were compared by calculating the ratio of normalized counts (UA^⬇^ counts / U^⬇^A counts). Ratios > 1 were scored as UA^⬇^ (RNase L) sites and ratios <1 were scored as U^⬇^A sites (EndoU) and total number of scored sites for either position are shown for each condition of viral infection in WT BMM at 9 and 12 hpi. (H) Model of EndoU and RNase L interaction at UA sites in MHV RNA.

Dinucleotide enrichment, a measurement comparing the frequency of cleavage at each dinucleotide to the frequency of each dinucleotide in the MHV genomic RNA, showed that U^⬇^A and C^⬇^A sequences were the only sequences with positively enriched cleavage in WT MHV-infected WT BMM (Fig. 4E, adjusted p-value (q) for fold change [log2(experiment / control)] of <1 x10^8***^). Dinucleotide enrichment and de-enrichment data for all dinucleotides at 9 and 12 hpi are available as supplemental data (Tables S1 and S2). These data indicate that EndoU cleaved MHV RNA at U^⬇^A and C^⬇^A sequences.

RNase L activity was also evident within MHV-infected WT BMM (Fig. 4A, B and C). RNase L activity, with characteristic cleavage predominantly after UA^⬇^ and UU^⬇^ dinucleotides, was significantly increased in both PDE^mut^-infected and EndoU^mut^-infected WT BMM (Fig. 4A). Dinucleotide enrichment showed that UA^⬇^, UU^⬇^ and UC^⬇^ sequences were positively enriched cleavage sites in PDE^mut^-infected and EndoU^mut^-infected WT BMM (Fig. 4B, adjusted p-value (q) for fold change [log2(experiment / control)] of <1×10^8***^). In IFNAR^-/-^ and RNase L^-/-^ BMM, the robust cleavage at UA^⬇^, UU^⬇^ and UC^⬇^ sequences decreased and pyrimidine specific cleavage dominated, especially in PDE^mut^-infected cells (Figs. S3A and S3B). These data indicate that RNase L cleaved MHV RNA after UA^⬇^, UU^⬇^ and UC^⬇^ sequences, consistent with other studies (31, 32).

The distinct specificity of cleavage for RNase L (UA^⬇^, UU^⬇^ and UC^⬇^ sequences) and EndoU (U^⬇^A and C^⬇^A sequences) allowed us to compare the relative amounts of each enzyme activity in the various experimental conditions. MHV RNAs were cleaved predominantly by EndoU activity within MHV^(S)^-infected and MHV^(V)^-infected BMM (Fig. 4A and D). MHV RNA was cleaved by both RNase L and EndoU activities within PDE^mut^-infected WT BMM while MHV RNA was cleaved predominantly by RNase L activity within EndoU^mut^-infected WT BMM (Fig. 4A and D). The activation of RNase L within PDE^mut^-infected and EndoU^mut^-infected WT BMM was expected, as these viral proteins coordinately block the OAS-RNase L pathway (22, 26, 28, 29). Dinucleotide analysis of positions downstream from cleavage sites confirmed a strong preference for adenine 3’ of the cleavage positions in MHV RNA in WT BMM (Figs. S2A and S2B). When EndoU was inactivated within EndoU^mut^-infected cells, the strong preference for adenine 3’ of cleavage positions in MHV RNA was dramatically reduced, but not entirely eliminated in WT BMM (Fig. S2A), IFNAR^-/-^ BMM (Fig. S3A) and RNase L^-/-^ BMM (Fig. S3B). The residual cleavage of MHV RNA within EndoU^mut^-infected RNase L^-/-^ BMM is likely due to angiogenin or another RNase A family member, as these enzymes are present within macrophage and they share a predilection for cleavage at U^⬇^A and C^⬇^A sequences (61–64).

We identified cyclic phosphate cDNAs dependent on the presence of either RNase L or EndoU and then used fold-change to identify and assign specific sites as RNase L or EndoU targets (Fig. 3 and Fig. S4). We determined how many of these sites could be assigned to either endoribonuclease for each experimental condition (Fig 3). EndoU cleaved MHV RNA at both 9 and 12 hpi in all three cell types, with increased amounts of cleavage at 12 hpi as compared to 9 hpi (Fig. 3A, B and C). MHV RNA was cleaved by RNase L activity at both 9 and 12 hpi in WT BMM, with exacerbated amounts of RNase L activity in PDE^mut^-infected and EndoU^mut^-infected WT BMM, as expected. In EndoU^mut^-infected WT BMM, there were nearly equal numbers of cleavage sites assigned to RNase L at 9 and 12 hpi, which was not observed in any other condition (Fig. 3A). By comparison with WT BMM, less RNase L-dependent cleavage was detected in IFNAR^-/-^ BMM (Fig. 3B), consistent with reduced OAS expression and reduced RNase L activity in IFNAR^-/-^ BMM (65). Additionally, the number of sites assigned to EndoU in IFNAR^-/-^ and RNase L^-/-^ BMM was less than that observed in WT BMM, suggesting that EndoU activity was altered in the absence of IFN signaling and innate immune effectors (Fig. 3A and 3B). We attributed the majority of endoribonuclease cleavage sites within MHV RNA to either EndoU (U^⬇^A and C^⬇^A sequences) or RNase L (UA^⬇^, UU^⬇^ and UC^⬇^ sequences) activities (Figs. 3 and 4); however, undefined enzymes cleaved MHV RNA within EndoU^mut^-infected RNase L^-/-^ BMM (Fig. 3C). As mentioned above, the residual cleavage of MHV RNA within EndoU^mut^-infected RNase L^-/-^ BMM was likely due to angiogenin or another RNase A family member, as these enzymes are present within macrophage and they share a predilection for cleavage at U^⬇^A and C^⬇^A sequences (61–64). The patterns and amounts of EndoU-dependent and RNase L-dependent cleavage in MHV RNA were consistent from one experiment (Figs. 3 and 4) to another (Figs. S8B-S8F).

It is intriguing to note that EndoU and RNase L share a common substrate dinucleotide, UA. Furthermore, we can distinguish between cleavage of UA by EndoU and RNase L as these enzymes cleave the UA sequence at distinct sites: EndoU cleaves between U^⬇^A sequences whereas RNase L cleaves after UA^⬇^ dinucleotides (Fig. 4H). We found hundreds of UA sequences in MHV RNA cleaved by both EndoU and RNase L (Fig. 4G). EndoU activity predominated in MHV^(S)^-infected and MHV^(V)^-infected WT BMM at 9 and 12 hpi (Fig. 4G, MHV^(S)^ and MHV^(V)^). Yet in PDE^mut^-infected WT BMM, either EndoU or RNase L cleaved about half of the UA sequences that were targeted by both enzymes (Figs. 4G and S8F, PDE^mut^). EndoU cleaved to a greater extent about half of the shared sites whereas RNase L cleaved another half to a greater extent (Figs. 4G and S8F, PDE^mut^). Thus, while EndoU and RNase L have overlapping sequence specificity and share common UA targets within MHV RNAs, these enzymes do not tend to cleave the same molecule at the same site at any one moment in time. Our data show that the majority of cleavage of MHV RNA was from EndoU rather than RNase L during WT MHV infections (Figs. 4G and S8F, WT); however, when the MHV PDE was mutated, a much larger proportion of cleavage events in viral RNA were from RNase L (Figs. 4G and S8F, PDE^mut^).

Taken together, these data indicate that EndoU and RNase L cleaved MHV RNA within infected BMMs. The majority of endoribonuclease cleavage sites within MHV RNA were attributed to either EndoU (U^⬇^A and C^⬇^A sequences) or RNase L (UA^⬇^, UU^⬇^ and UC^⬇^ sequences) activities (Figs. 3 and 4). However, data from EndoU^mut^-infected RNase L^-/-^ BMM (Fig. 3C) indicate that viral RNA was cleaved by other undefined endoribonucleases as well. Furthermore, when MHV NS2 PDE or nsp15 EndoU were inactivated by mutations, RNase L activity was much greater, with increased cleavage of MHV RNA by RNase L. Thus, both MHV NS2 PDE and nsp15 EndoU activities prevent MHV RNA cleavage by the dsRNA-activated OAS/RNase L pathway, confirming our previous reports (26, 28, 29).

### RNase L-dependent and EndoU-dependent cleavage sites in MHV RNA

A fold-change analysis was used to compare the magnitudes of RNase L-dependent and EndoU-dependent cleavage at each base of MHV RNA across experimental conditions (Fig. 5A). By subtracting endoribonuclease cleavage events detected for each virus in RNase L^-/-^ BMM, we identified the top 100 RNase L-dependent cleavage sites in MHV RNA (Fig. 5B). By subtracting the endoribonuclease cleavage events detected for the EndoU^mut^, we identified the top 100 EndoU-dependent cleavage sites in MHV RNA (Fig. 5C).

**Figure 5.**
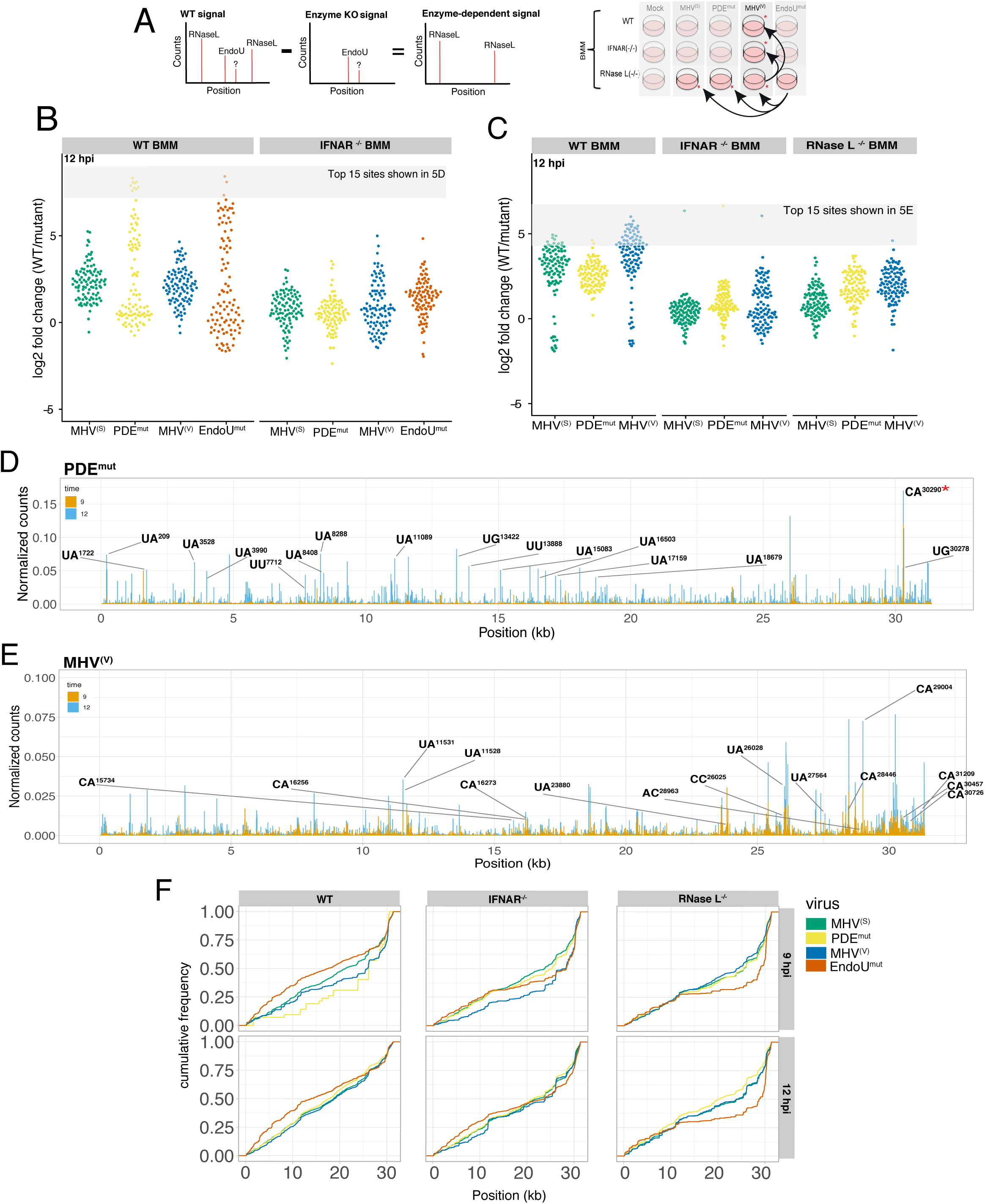
RNase L-dependent and EndoU-dependent cleavage sites in MHV RNA. (A) Schematic outline of analysis to identify EndoU/RNase L-dependent cyclic phosphate reads. (B and C) Fold change values for the top 100 RNase L-dependent or EndoU-dependent cleavage sites. Fold change in cyclic phosphate signal when comparing WT or IFNAR^(-/-)^ BMM infected with MHV^(S),^ MHV^(V)^, PDE^mut^, and EndoU^mut^ virus to RNase L^-/-^ BMM (B) or MHV^(S),^ MHV^(V)^, PDE^mut^ virus to infection with EndoU^mut^ virus across all cell types (C) displayed as violin scatter plot. Log2 fold change in the absence of RNase L activity (B) or in the absence of EndoU activity (C) was calculated for each position in the MHV genomic RNA. Fold change values for the top 100 RNase L-dependent or EndoU-dependent sites were compared in WT and IFNAR^-/-^ BMM under conditions of infection with MHV^(S),^ MHV^(V)^, PDE^mut^, and EndoU^mut^ virus at 12 hpi (B) or in all cell types across conditions of infection with MHV^(S),^ MHV^(V)^, PDE^mut^ virus at 12 hpi. (D) Frequency and location of RNase L-dependent cleavage sites in MHV RNA. Cyclic phosphate counts at each position in the viral genome were normalized by removing signal that occurred in the absence of RNase L, which emphasizes sites that are RNase L-dependent in WT BMM infected with MHV^(S)^, and PDE^mut^ at 9 and 12 hpi. Labeled positions and dinucleotides (−2 base : −1 base) on the graph of PDE^mut^ represent the top 15 RNase L-dependent cleavage sites (B) with the greatest fold-change in RNase L activity (*site with robust cleavage without canonical RNase L dinucleotide preference and independent of EndoU activity; not identified as top site by RNase L fold change analysis). (E) Frequency and location of EndoU-dependent cleavage sites in MHV RNA. Cyclic phosphate counts at each position in the viral genome were normalized by removing signal that occurred in the absence of EndoU, which emphasizes sites that are EndoU-dependent and RNase L-independent in RNase L^-/-^ BMM infected with WT MHV^(V)^ at 9 and 12 hpi. Labeled positions and dinucleotides (−1 base : +1 base) represent the top 15 EndoU-dependent cleavage sites with the greatest fold-change in EndoU activity (C). (F) Cumulative distribution of normalized counts by position of MHV genome for every position with >= 10 cyclic phosphate counts across all cell types and infection conditions.

RNase L-dependent sites in MHV RNA were cleaved at the greatest magnitudes in PDE^mut^-infected and EndoU^mut^-infected WT BMM (Fig. 5B). RNase L-dependent cleavage of MHV RNA was substantially lower in IFNAR^-/-^ cells, as expected (65), especially that associated with infections by the PDE^mut^ and EndoU^mut^ (Fig. 5B). The top 15 RNase L-dependent cleavage sites in MHV RNA were at UA^⬇^, UU^⬇^ and UG^⬇^ dinucleotides distributed across the viral genome, with a clustering of sites within the first 2/3 of the genome (Fig. 5D). Magnitudes of cleavage at each of these sites ranged from 0.05 to 0.08% of all cleavage sites in each cDNA library (∼1/2000 cleavage sites in the cDNA library). Together, these top 15 cleavage sites in MHV RNA accounted for ∼1% of all cleavage sites in this cDNA library, across all host and viral RNAs.

These data indicate that RNase L cleaved coronavirus RNA most efficiently at a relatively small number of sites in the viral genome.

EndoU-dependent cleavage sites in MHV RNA were evident in WT, IFNAR^-/-^ and RNase L^-/-^ BMMs; however, EndoU cleaved MHV RNA to a greater extent in WT BMM (Fig. 5C). Subdued magnitudes of EndoU-dependent cleavage of MHV RNA were observed at 12 hpi in IFNAR^-/-^ and RNase L^-/-^ cells, as compared to WT BMM, suggesting a potential functional interaction between EndoU and dsRNA-activated host responses, or RNase L in particular. Additionally, most of the sites with EndoU-dependent cleavage activity had similar magnitudes of change, leading to a uniform distribution of sites across all conditions, excluding a few outliers. The top 15 EndoU-dependent cleavage sites in MHV RNA were at C^⬇^A and U^⬇^A sequences distributed to a greater extent in the last 2/3 of the viral genome (Fig. 5E).

We examined the cumulative distribution of cleavage in MHV RNA, across all conditions (Figs. 5F and S8E). In this analysis, we plotted the overall accumulation of cyclic phosphate reads as a function of position along the MHV genomic RNA (Figs. 5F and S8E). Because RNase L-dependent cleavage sites (Fig. 5D) and EndoU-dependent cleavage sites (Fig. 5E) were distributed across the MHV RNA genome in WT BMM, cumulative cleavage increased from 0% at the 5’ end of the genome to 100% at the 3’ end of the genome, with a slope of ∼45° for MHV^(S)^ and MHV^(V)^ in WT BMM [Fig. 5F, WT BMM, green and blue lines for MHV^(S)^ and MHV^(V)^]. In EndoU^mut^-infected WT BMM, cleavage of MHV RNA increased in the ORF1a and ORF1b regions of the genome as compared to MHV^(S)^ and MHV^(V)^, shifting the slope of cumulative cleavage to the left (Fig. 5F, WT BMM, red line for EndoU^mut^). In contrast, when both EndoU and RNase L activities were absent, as in in EndoU^mut^-infected RNase L^-/-^ BMM, cleavage of MHV RNA was substantially reduced across most of the genome, with a spike of EndoU- and RNase L-independent cleavage near the 3’ UTR (Fig. 5F, RNase L^-/-^ BMM, red line for EndoU^mut^). Note how the slope of the line for EndoU^mut^ goes from ∼50% to 100% of cumulative cleavage between nucleotides 30,000 and 31,344. This indicates that endoribonucleolytic cleavage was much more pronounced near the 3’ terminus of MHV RNA in EndoU^mut^-infected IFNAR^-/-^ and RNase L^-/-^ BMM, as compared to WT BMM. These data indicate that EndoU^mut^ MHV RNA was cleaved at very different magnitudes from one end to the other in WT BMM versus that in RNase L^-/-^ BMM, with increased relative amounts of cleavage between nts 1-20,000 in WT BMM, less cleavage between nts 1-30,000 in RNase L^-/-^ BMM, and a spike in cumulative cleavage near the 3’ terminus in RNase L^-/-^ BMM.

These data also indicate that EndoU and RNase L account for a substantial amount of the cumulative cleavage in the orf1a and orf1b regions of the MHV RNA genome. MHV RNA was cleaved to a greater extent within orf1a and orf1b in WT BMM, especially when EndoU was disabled (Fig. 5F, red line for EndoU^mut^ shifts to the left in WT BMM). Conversely, MHV RNA was cleaved to a lower extent within orf1a and orf1b in RNase L^-/-^ BMM, especially when EndoU was disabled (Fig. 5F, red line for EndoU^mut^ shifts to the right in RNase L^-/-^ BMM). When EndoU and RNase L activities were absent, as in EndoU^mut^-infected RNase L^-/-^ BMM, the residual cleavage of MHV RNA by unspecified endoribonucleases occurred predominantly near the 3’ terminus of the viral genome.

### Endoribonuclease cleavage sites in distinct MHV RNA sequences and structures

We next examined the frequency of endoribonuclease cleavage in distinct regions of MHV RNA (Figs. 6 and S7). The cumulative amounts of cleavage in each region of MHV RNA were plotted unadjusted (Fig. 6A) or adjusted for both RNA abundance and size (Fig. 6B). In supplemental data we show cleavage adjusted for RNA abundance alone (Fig. S7A) or size alone (Fig. S7B). Cleavage was detected in every region of MHV RNA, from the 5′ NTR to the 3′ NTR, including relatively small TRS sequences (Figs. 6A and 6B). The vast majority of cleavage events occurred in 1a/1b, S and N open reading frames (Fig. 6A). When adjusted for MHV RNA abundance, cleavage was most frequent in the ORF 1a/1b region and the ns2, HE and S ORFs (S7A). Furthermore, with adjustments for size and abundance (Figs. 6B), one can see that each of the TRS elements was targeted for cleavage at frequencies similar to that observed in Orf1a/1b. Thus, although TRS sequences are quite small, they were cleaved just as frequently as RNA sequences in other regions of MHV RNA. Intriguingly, TRS6 was targeted more frequently (by EndoU) than other regions of MHV RNA, including other TRS elements (Fig. 6C). TRS6, with a UCCAAAC sequence, is distinct from other TRS elements, which possess UCUAAAC sequences. We detected the most robust EndoU-dependent cleavages at C^⬇^A and U^⬇^A dinucleotides of TRS elements 4, 6, and 7 (Fig. 6C). In TRS elements 4 and 6, cleavage at the very 3’-end of the TRS sequence was dependent on the presence of a downstream adenine outside of the TRS sequence (Fig. 6C). Interestingly, the upstream C^⬇^A cleavage site in TRS 6 (Fig. 6C) relies on one of the single nucleotide polymorphisms (28960 T > C) that we detected in the viral genomes (Table S3). In vitro studies using purified EndoU show cleavage of a U^⬇^A dinucleotide within a TRS substrate (23). Our data indicate that C^⬇^A and U^⬇^A dinucleotides of TRS elements are physiologic targets of EndoU.

**Figure 6.**
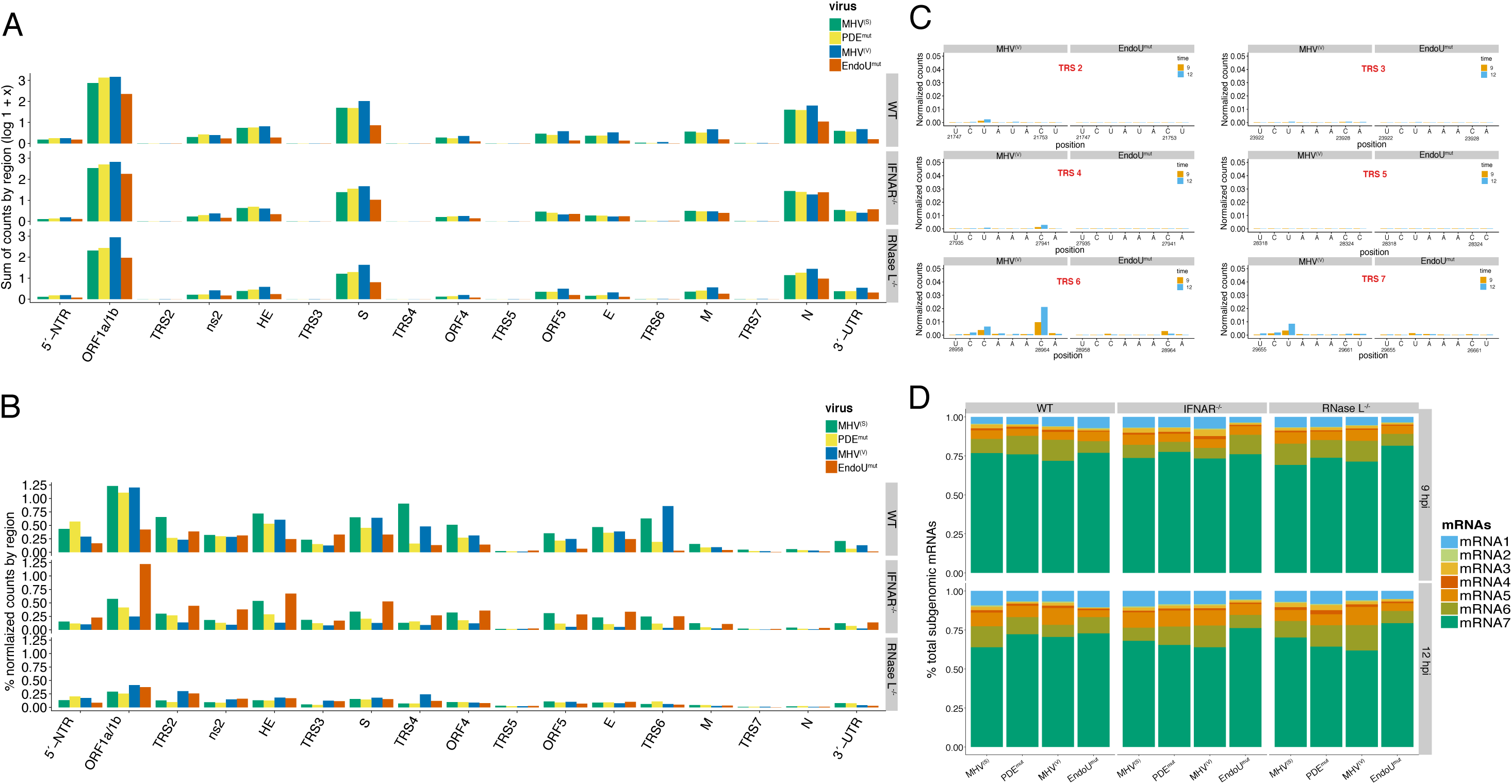
Abundance of cyclic phosphate ends by MHV genomic region and MHV mRNA abundance. Sum of endonuclease cleavage sites in MHV RNA, by genomic regions: sum of cyclic phosphate reads (A), sum of cyclic phosphate reads normalized by MHV mRNA abundance (B) or sum of cyclic phosphate reads normalized by the length of the MHV genomic region (C). (A) Sum of cyclic phosphate cDNA reads displayed by MHV RNA region for WT, IFNAR^-/-^, RNase L^-/-^ BMM across all conditions of viral infection at 12 hpi. Transcriptional regulatory sequences (TRS) are numbered by their associated mRNA (2–7). Other MHV genomic regions are labeled as shown in Figure 1A. (B) Frequency of endonuclease cleavage sites in MHV RNA, by genomic regions, normalized by MHV mRNA abundance. Sum of cyclic phosphate counts normalized by mRNA abundance at each capture base, displayed by MHV genomic region for WT, IFNAR^-/-^, RNase L^-/-^ BMM across all conditions of viral infection at 12 hpi. (C) Percent of sum of normalized counts per length of MHV genomic region for WT, IFNAR^-/-^ and RNase L^-/-^ BMM across all conditions of viral infection at 12 hpi. Dotted line represents baseline percent of cleavage expected by cell type ([total number of cyclic phosphate counts / total genome size x 100]). (D) Frequency and location of cleavage in the MHV TRS elements in WT BMM during infection with MHV^(V)^ and EndoU^mut^ at 12 hpi. The x-axis includes the sequence and position of the 6-base MHV TRS elements. (E) Normalized counts (sum of MHV sg mRNA / sum of all MHV mRNAs) of MHV sg mRNAs detected in WT, IFNAR^-/-^, RNase L^-/-^ BMM across all conditions of viral infection at 9 and 12 hpi. (F) Sum of all MHV sg mRNAs (RPM) for WT, IFNAR^-/-^, RNase L^-/-^ BMM across all conditions of viral infection at 9 and 12 hpi.

RNAseq was used to measure the abundance of MHV RNA in all experimental conditions (Figs. S5 and 6D). MHV RNA was abundant in all samples from virus-infected cells, with similar amounts of MHV RNA across conditions, but for EndoU^mut^-infected WT BMM at 9 and 12 hpi (Fig. S5A). Decreased amounts of EndoU^mut^ RNA in WT BMM (Fig. S5A) correlated with decreased virus replication in EndoU^mut^-infected WT BMM at 9 and 12 hpi (26). RNAseq reads were detected across the MHV RNA genome, with the most abundant reads corresponding to leader sequences at the 5’ end of the genome and sg mRNA sequences at the 3’ end of the genome (Fig. S5B). Consistent with published studies (53), MHV mRNA7 was most abundant, accounting for 70 to 80% of MHV mRNAs (Figs. 6D and S9C). MHV mRNAs 1-7 were present in all conditions, with some changes in relative amounts from one condition to another (Figs. 6D and S9C). MHV mRNA1 (genomic RNA) was increased proportionally to other MHV mRNAs in EndoU^mut^-infected WT BMM at 12 hpi. MHV mRNA 7 was increased relative to other MHV mRNAs at 12 hpi in EndoU^mut^-infected IFNAR^-/-^ BMM and RNase L^-/-^ BMM. Remarkably, MHV RNA abundance did not correlate with the frequency of cyclic phosphate reads in viral RNA (Figs. S11A & S11B). Altogether, these data indicate that MHV RNA replication was able to produce each of the MHV mRNAs in proportional amounts, despite considerable changes in endoribonuclease activity from one condition to another.

Endoribonuclease cleavage sites were detected in functional RNA sequences and structures, including the Orf1a/1b frameshift element and the MHV 3′ NTR (Fig. 7). The Orf1a/1b frameshift element contains both RNase L-dependent and EndoU-dependent cleavage sites (Figs. 7A and 7B). Likewise, the MHV 3′ NTR contains both RNase L-dependent and EndoU-dependent cleavage sites (Fig. 7C). The MHV 3′ NTR spans nucleotide 31034, adjacent to the N stop codon, to nucleotide 31334, adjacent to the poly(A) tail (Fig. 7C). Functional RNA sequences and structures within the 3′ NTR include an essential bulged stem-loop (nts 31034-31100), an essential pseudoknot (nts 31101-31150), a non-essential hypervariable region (HVR) (nts 31179-31288), a polyadenylation signal (nts 31293-31298) and a poly(A) tail (66–68). A number of EndoU-dependent cleavage sites were detected within the 3′ NTR, including prominent cleavage sites immediately adjacent to the poly(A) tail (Fig. 7C, ^31332^C^⬇^AC^⬇^A^31335^). Together, these two cleavage sites account for ∼0.15% of all cleavage sites in the cDNA library for the WT MHV in WT BMM at 12 hpi, corresponding to ∼1/677 cleavage sites in the entire cDNA library.

**Figure 7.**
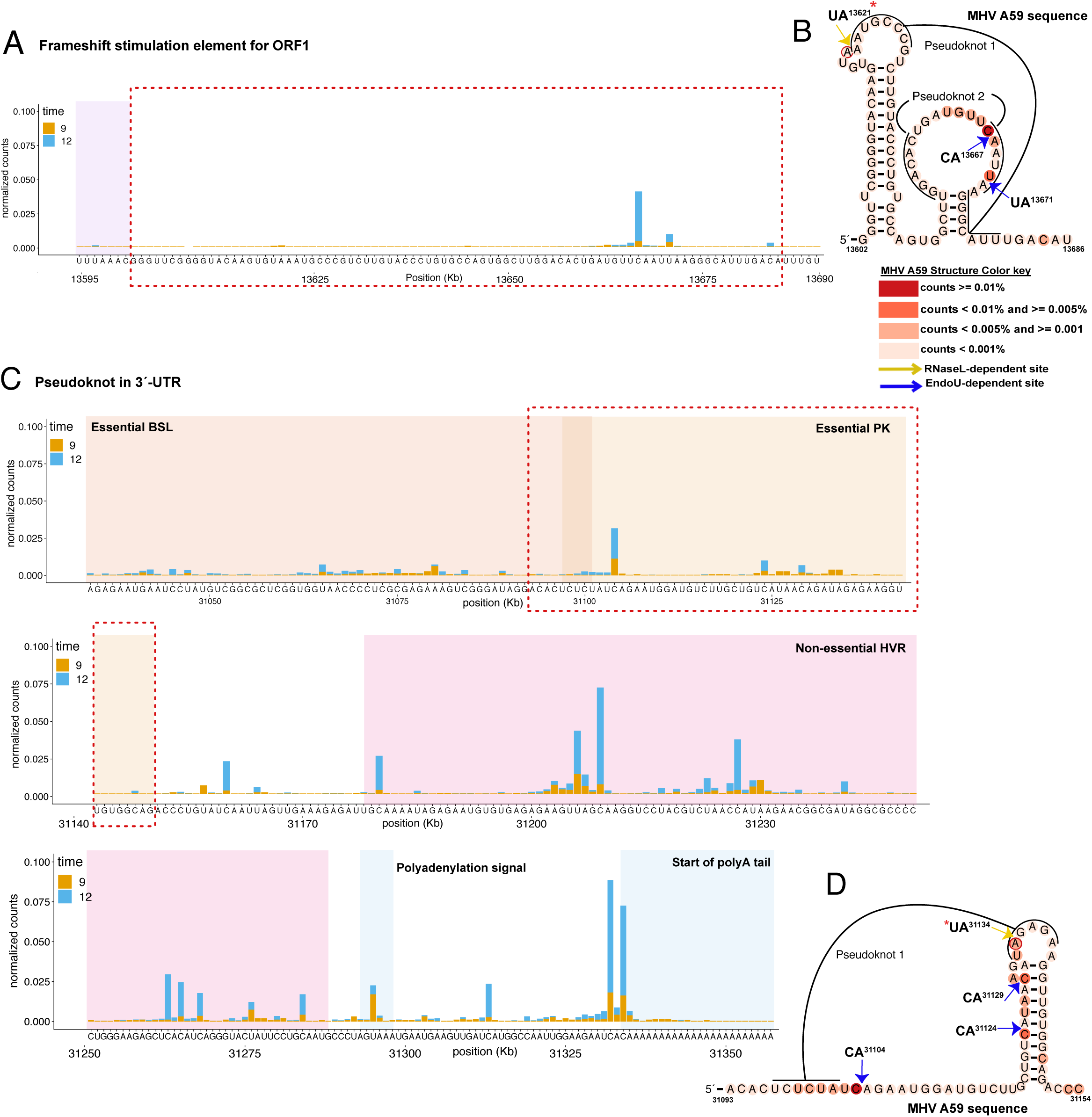
MHV secondary structures associated with RNase L-dependent and EndoU-dependent cleavage sites. (A and C) Nucleotide resolution graphs displaying normalized counts by position for the regions encompassing secondary structure predictions. (B and D) Secondary structures of frameshift stimulation element (B) and MHV 3’-UTR pseudoknot (D), generated using available consensus alignment and the R-scape program (85). MHV A59 sequence mapped to consensus secondary structures using available covariation model and the Infernal program (86). Base coloring of MHV A59 sequence based on normalized cDNA reads as indicated in key for 12 hpi in WT BMM infected with MHV^(V)^. *Base RNase L-dependent cleavage activity is increased in PDE^mut^ or EndoU^mut^ infection as compared to MHV^(V)^ infection.

When EndoU was inactivated by an H277A mutation, the cleavage of MHV RNA at the ^31332^C^⬇^AC^⬇^A^31335^ sequences adjacent to the poly(A) tail was dramatically reduced, but not entirely eliminated, in WT BMM (Fig. S6A). Furthermore, there was EndoU-independent cleavage of MHV RNA at the ^31332^C^⬇^AC^⬇^A^31335^ sequence in IFNAR^-/-^ BMM (Fig. S6B) and RNase L^-/-^ BMM (Fig. S6C). Cleavage of MHV RNA at the ^31332^C^⬇^AC^⬇^A^31335^ sequences adjacent to the poly(A) tail were notable whether unadjusted (Fig. S6A-C) or adjusted for RNA abundance (Fig. S6D-F). These data indicate that the ^31332^C^⬇^AC^⬇^A^31335^ sequence in MHV RNA was susceptible to both EndoU-dependent and EndoU-independent cleavage. The EndoU-dependent cleavage of the ^31332^C^⬇^AC^⬇^A^31335^ sequence in MHV RNA was substantially greater than the EndoU-independent cleavage in WT BMM (Fig. S6A); however, substantial amounts of EndoU-independent cleavage were detected at the ^31332^C^⬇^AC^⬇^A^31335^ sequence in IFNAR^-/-^ BMM (Fig. S6B) and RNase L^-/-^ BMM (Fig. S6C).

### Cleavage of rRNA and changes in host gene expression

Because RNase L cleaves 18S rRNA at specific sites in human cells (31, 32), we examined RNase L-dependent cleavage of 18S rRNA within MHV-infected murine BMMs (Figs. 8A, 8B and S10). A fold-change analysis was used to compare the magnitudes of RNase L-dependent cleavage at each base of 18S rRNA across experimental conditions (Fig. 8A). By subtracting endoribonuclease cleavage events detected for each virus in RNase L^-/-^ BMM, we identified the top 100 potential RNase L-dependent cleavage sites in MHV RNA (Fig. 8B). Four RNase L-dependent cleavage sites were clearly evident in 18S rRNA: UU^542^, UU^543^, UU^771^ and UA^772^. These sites, on the surface of 18S ribosomal subunits, are analogous to RNase L-dependent cleavage sites in human 18S subunits (31, 32). 18S rRNA was cleaved at these sites to a significant magnitude in PDE^mut^-infected and EndoU^mut^-infected WT BMM (Figs. 8A, 8B and S10). 18S rRNA was not cleaved at significant magnitudes within mock-infected BMM nor in IFNAR^-/-^ or RNase L^-/-^ BMM (Fig. 8A and 8B). Thus, as in human cells (31, 32), RNase L targets 18S rRNA for cleavage at precise sites in murine cells. Furthermore, RNase L activity was specifically increased within PDE^mut^-infected and EndoU^mut^-infected WT BMM, as compared to MHV^(S)^-infected and MHV^(V)^-infected WT BMM (Fig. 8A and 8B). Although RNase L-dependent cleavage sites in rRNA were easily detected (Figs. 8A, 8B and S10), EndoU-dependent cleavage sites in rRNA were not detected (Fig. S10B). These data show the dsRNA-dependent OAS/RNase L pathway was significantly activated in PDE^mut^- and EndoU^mut^-infected WT BMM, and exclude rRNAs as targets of EndoU. dsRNA-dependent host gene expression was also increased within MHV-infected WT BMM (Fig. 8C-E). We used a Volcano plot (Fig. 8C) and gene ontology (GO) analyses to classify the nature of host gene expression within MHV-infected WT BMM (Fig. 8D and 8E) (49, 50). The Volcano plot shows the expression of many genes increasing by 2^2^ to 2^10^-fold / 4-fold to 1024-fold (Fig. 8C). Increased gene expression in MHV^(S)^-infected WT BMM corresponded to a number of biological processes: negative regulation of apoptosis, LPS-activated gene expression, positive regulation of GTPase activity and IFN-gamma activated gene expression (Fig. 8D). Increased gene expression in EndoU^mut^-infected WT BMM (Fig. S12A and S12B) corresponded to some of these same groups of host genes, with a notable addition, response to exogenous dsRNA (Fig. 8E). Thus, GO analysis indicated that host gene expression associated with response to exogenous dsRNA was specifically activated in EndoU^mut^-infected WT BMM, as compared to MHV^(S)^-infected WT BMM (Fig. 8D and 8E).

**Figure 8.**
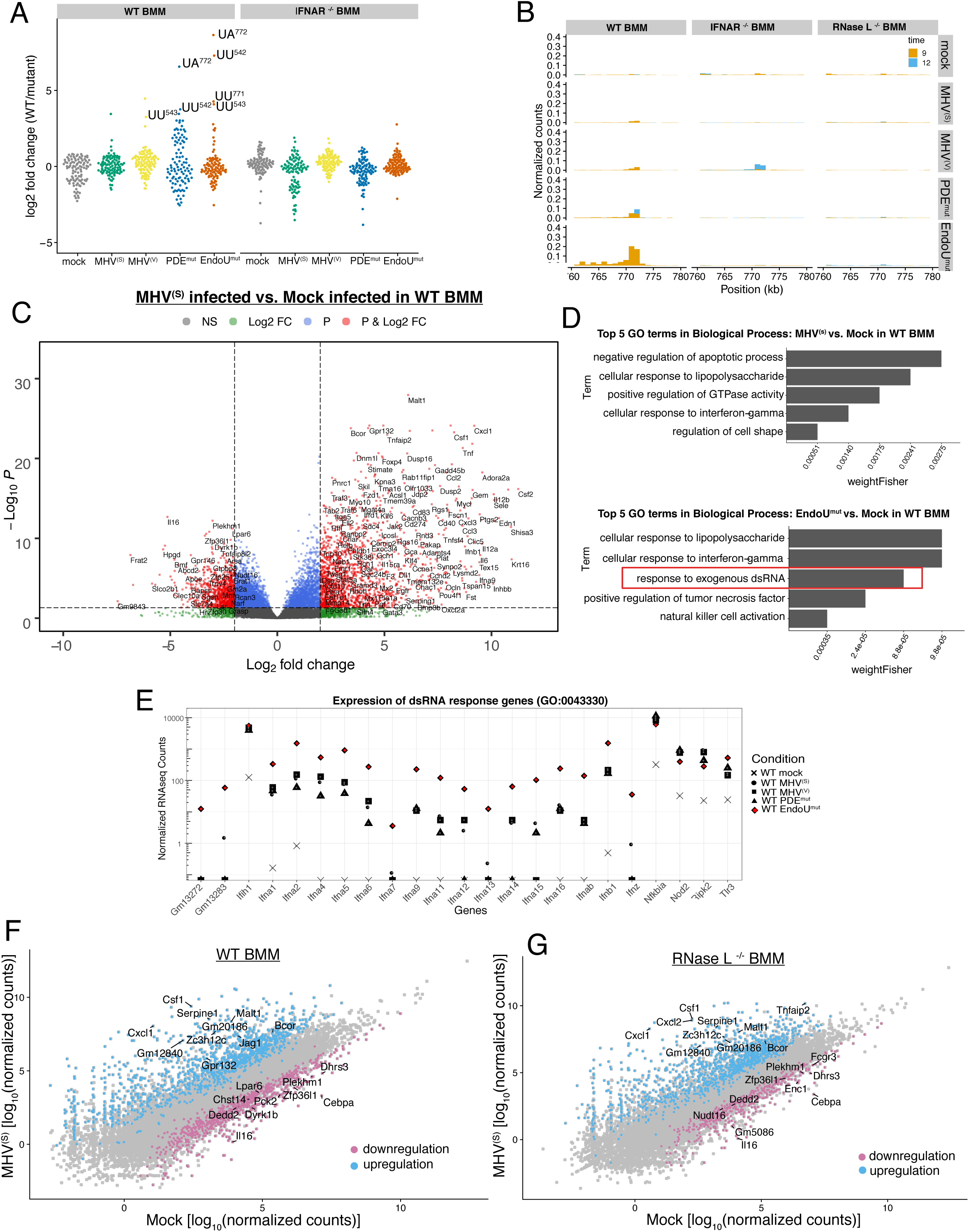
Endoribonuclease cleavage of cellular RNAs and changes in host gene expression. RNase L-dependent cleavage sites in 18S rRNA (A & B). (A) RNase L-dependent cleavage sites in 18S rRNA by fold change in signal when comparing WT or IFNAR^-/-^ BMM mock-infected or infected with MHV^(S),^ MHV^(V)^, PDE^mut^, and EndoU^mut^ virus to RNase L^-/-^ BMM. Log2 fold change in the absence of RNase L activity was calculated for each position in the rRNA. The distribution of the top 100 RNase L-dependent cleavage sites were compared in WT and IFNAR^-/-^ BMM under conditions of mock infection or infection with MHV^(S),^ MHV^(V)^, PDE^mut^, and EndoU^mut^ virus at 9 hpi. 18S rRNA was cleaved in an RNase L-dependent manner at UU^542^, UU^543^, UU^771^ and UA^772^. (B) RNase L-dependent cleavage of 18S rRNA at UU^771^ and UA^772^ at 9 and 12 hpi, predominantly in MHV PDE^mut^- and EndoU^mut^-infected WT BMM. (C) Volcano plot of changes in host gene expression comparing MHV^(s)^-infected and mock-infected WT BMM. Host genes differentially expressed (FDR < 0.05) and upregulated (logFC > 2) or downregulated (logFC < −2). (D) GO analysis: MHV EndoU^mut^ infection of WT BMM provokes increased expression of host genes associated with exogenous dsRNA response. Categories of biological processes with significantly upregulated genes (p < 0.01, lo2FC > 2) identified by comparing MHV^(s)^-infected and EndoU^mut^-infected WT BMM to mock-infected WT BMM. Top 5 categories significantly enriched (weightFisher < 0.01). (E) Expression of host genes in GO category “response to exogenous dsRNA”. Expression (log_10_ normalized counts) of genes in the GO category “response to exogenous dsRNA” for WT BMM at 12 hpi: mock-infected (◼) or MHV-infected with MHV^(S)^ (●), MHV^(V)^ (▲), PDE^mut^ (+), and EndoU^mut^ (red-circle in black square). (F and G) Differential host gene expression comparing mock-infected and MHV^(s)^-infected cells at 12 hpi: WT BMM (F) and RNase L^-/-^ BMM (G). Upregulated (fold change > 2, FDR < 0.01) and downregulated transcripts (fold change < −2, FDR < 0.01).

Because GO analysis implicated “response to exogenous dsRNA”, we examined the magnitudes of expression for each host gene in this gene ontology group: GM13272, GM13283, IFN-alpha genes, IFN-beta, IFN-Z, Nfkbia, Nod2, Ripk2 and Tlr3 (Fig. 8E). We compared magnitudes of expression in mock-infected, MHV^(S)^-infected, MHV^(V)^-infected, PDE^mut^-infected and EndoU^mut^-infected WT BMM (Fig. 8E). Host gene expression associated with response to dsRNA increased by 100- to 1000-fold in MHV-infected WT BMM as compared to mock-infected cells, with even larger 1000- to 10,000-fold increases in EndoU^mut^-infected WT BMM (Fig. 8E). Thus, host genes associated with response to dsRNA were notably increased in MHV-infected BMM, with the greatest increases occurring within EndoU^mut^-infected WT BMM (Fig. 8E). Genes upregulated or downregulated in MHV^(S)^-infected cells did not vary substantially between WT and RNase L^-/-^ BMM (Fig. 8F and 8G). Altogether, these data indicate that dsRNA-dependent host responses were exacerbated within MHV-infected cells, especially in EndoU^mut^-infected WT BMM. These data are consistent with recent studies from the Baker lab (69).

### Cellular endoribonucleases

The residual cleavage of MHV RNA within EndoU^mut^-infected RNase L^-/-^ BMM provoked our consideration of other cellular endoribonucleases. We hypothesized that residual pyrimidine specific cleavage of MHV RNA might be due to one or another RNase A family enzyme (63). We also considered T2 endoribonucleases based on their reported contributions to TLR8 activation (70). Consequently, we examined the expression of RNases 4 and 5 (angiogenin) and RNases T2A and T2B (Fig. S12C). Changes in magnitudes of RNase 4 and 5 expression were observed, with ∼10-fold decreased expression in MHV^(S)^-infected and MHV^(V)^-infected WT BMM as compared to Mock-infected WT BMM (Fig. S12C). Decreased expression of RNases 4 and 5 was not as strong in PDE^mut^-infected WT BMM, and very little decrease in expression was observed in EndoU^mut^-infected WT BMM. Similar changes in expression of RNases 4 and 5 were observed in IFNAR^-/-^ BMM and RNase L^-/-^ BMM, with significantly decreased expression in MHV^(S)^-infected and MHV^(V)^-infected cells and a more limited decrease in EndoU^mut^-infected cells (Fig. S12C). Because RNases 4 and 5 share a complex dual promoter (71), with alternative splicing leading to the expression of either RNase 4 or RNase 5, coordinate increases and decreases in their expression was not unexpected. These data reinforce our suspicion regarding the residual pyrimidine specific cleavage of MHV RNA within EndoU^mut^-infected RNase L^-/-^ BMM.

In contrast to expression of RNase 4 and 5, changes in magnitudes of expression of RNases T2A and T2B were relatively small within MHV-infected cells, with a tendency for slightly increased expression (Fig. S12C). RNase T2 cleaves RNA within endosomes and lysosomes, targeting purine:uridine dinucleotides, R^⬇^U (70). The residual purine specific cleavage of MHV RNA within EndoU^mut^-infected RNase L^-/-^ BMM might be associated with RNase T2 activity; however, our experiments do not definitively address this possibility.

## DISCUSSION

We address a key question in the coronavirus field (72): What is the natural target of EndoU? Coronavirus EndoU prevents dsRNA-activated antiviral responses in infected cells (26); however, it is not clear how EndoU does this because its physiologic RNA substrates are unknown. In this study, we used MHV-infected bone marrow-derived macrophage (BMM) and cyclic phosphate cDNA sequencing to identify the RNA targets of EndoU.

We found that EndoU targeted MHV RNA within infected cells, cleaving viral RNA on the 3′ side of pyrimidines with a strong preference for cleavage between U^⬇^A and C^⬇^A sequences (endoY^⬇^A) (Fig. 4). This cleavage specificity from MHV-infected cells is consistent with that of purified EndoU (23, 60) and RNase A (61, 62), enzymes that are functionally and structurally related to one another (7). EndoU cleavage was detected in every region of MHV RNA, from the 5′ NTR to the 3′ NTR, including relatively small TRS sequences (Figs. 5E, 6 and 7). Because MHV RNA is a template for both viral mRNA translation and viral RNA replication, cleavage by EndoU could inhibit both of these biosynthetic processes (Fig. 9). Intriguingly, MHV TRS sequences contain EndoU target sequences (C^⬇^A and U^⬇^A sequences) (Fig. 6C). TRS6, which was targeted more frequently by EndoU than other TRS elements, contains a C^⬇^A target sequence rather than a U^⬇^A sequence. We postulate that EndoU cleaves MHV RNA in a regulated manner, to inhibit negative-strand RNA synthesis, thereby preventing the accumulation of viral dsRNA (Fig. 9). Nsp16 (2′ O-MT) could regulate EndoU-mediated cleavage of MHV RNA by methylating C^⬇^A and U^⬇^A sequences (1).

**Figure 9.**
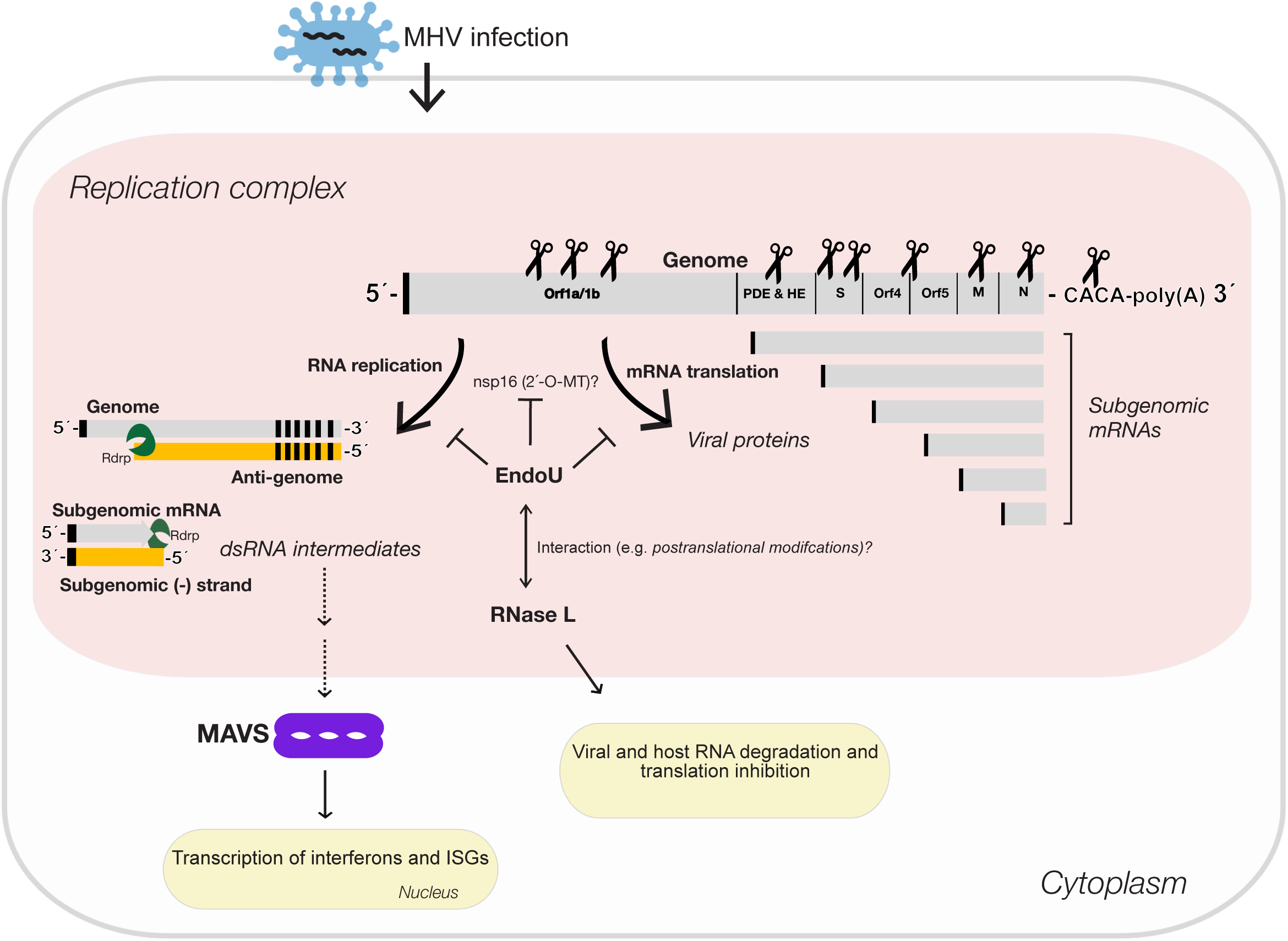
EndoU targets in MHV RNA. MHV RNA was targeted for cleavage by EndoU within infected BMM. MHV RNA was cleaved by EndoU in all regions of the genome, at C^⬇^A and U^⬇^A sequences. Because MHV RNA is a template for both viral mRNA translation and viral RNA replication, cleavage by EndoU could inhibit both of these biosynthetic processes. Intriguingly, MHV TRS sequences contain EndoU target sequences (C^⬇^A and U^⬇^A sequences). TRS6, which was targeted more frequently by EndoU than other TRS elements, contains a C^⬇^A target sequence rather than a U^⬇^A sequence. We postulate that EndoU cleaves MHV RNA in a regulated manner, to inhibit negative-strand RNA synthesis, thereby inhibiting the accumulation of viral dsRNA. Nsp16 (2’ O-MT) could regulate EndoU-mediated cleavage of MHV RNA by methylating C^⬇^A and U^⬇^A sequences. EndoU and RNase L cleave an overlapping set of UA sequences within MHV, suggesting a functional interplay between host and viral endoribonucleases.

### How does EndoU inhibit double-stranded RNA-activated antiviral responses?

Coronavirus EndoU prevents the activation of multiple host dsRNA sensors, including MDA5, OAS and PKR (22, 26, 27). dsRNA-activated OAS/RNase L and PKR pathways restrict the replication of EndoU-deficient coronaviruses (26). Because EndoU^mut^-infected cells had increased accumulation of dsRNA, Kindler and colleagues (26) concluded that EndoU functions as a viral RNA decay pathway to evade dsRNA-activated antiviral host cell responses. Consistent with this idea, Hackbart et al. (73) report that EndoU targets poly(U) sequences at the 5’ end of viral negative-strand RNA. Another report suggests that EndoU might control the localization of viral dsRNA within cells, perhaps maintaining dsRNA within membranous RNA replication complexes (22). Our data suggest a third possibility, that EndoU targets MHV RNA to prevent the synthesis of dsRNA (Fig. 9): EndoU-dependent cleavages were detected throughout the genomic RNA (Figs. 5E, 6 and 7), indicating that EndoU destroys the template for negative-strand RNA synthesis, precluding the formation of dsRNA, rather than acting on dsRNA. Cyclic phosphate cDNA sequencing detected large amounts of cleavage in MHV (+) strand (Fig. 2 and Fig. 8SA) and vanishing little cleavage in MHV (-) strand (Figs. S1B, S11C-D). Cyclic phosphate cDNA sequencing can readily detect cleavage sites in both (+) and (-) strands of viral RNA (31, 32); however, cleavage of poly(U) sequences at the 5’ end of MHV negative-strand RNA cannot be detected because the resulting cyclic phosphate RNA fragments are too small (<20 bases long) and they are homopolymeric, preventing detection by our sequencing and bioinformatics pipelines. While it is possible that EndoU targets poly(U) sequences, the specificity of EndoU for C^⬇^A and U^⬇^A sequences in vivo (Fig. 4) is inconsistent with poly(U) substrates being physiologically relevant. Furthermore, purified EndoU (60) and RNase A (61, 62) readily target UA sequences within heteropolymeric substrates. Thus, we conclude that EndoU targets MHV (+) strand RNA to prevent the synthesis of dsRNA (Fig. 9). Nonetheless, potential RNA substrates in (+) and (-) strands are not mutually exclusive. EndoU-dependent cleavage of the CACA sequences at the 3’ end of the (+) strand and the poly(U) at the 5’ end of the (-) strand could occur coordinately, as both are co-localized adjacent to one another at the same end of dsRNA products. When EndoU was mutated, we detected the activation of the dsRNA-dependent OAS/RNase L pathway (Figs. 4, 5 and 8) and increased host gene expression associated with response to dsRNA (Figs. 8D and 8E). These data, like other reports (22, 26, 27, 69), indicate EndoU prevents the activation of dsRNA sensors.

EndoU cleaved MHV RNA in every region of the genome (Figs. 5E, 6 & 7). Because MHV RNA is a template for both viral mRNA translation and viral RNA replication, cleavage by EndoU could inhibit both of these biosynthetic processes (Fig. 9). Cleavage of the viral genome would prevent the expression of the viral replicase. Coronavirus RNA synthesis requires ongoing expression of the viral replicase, with negative-strand RNA synthesis being most dependent on new replicase expression (74). Substantial amounts of EndoU-dependent cleavage were detected in orfs 1a and 1b, especially within WT BMM (Fig. 5), potentially limiting the expression of replicase. Cleavage of MHV genomic RNA, the template for both genomic and subgenomic negative-strand RNA synthesis (10), would also prevent the synthesis of dsRNA products (Fig. 9). EndoU-mediated cleavage of the tandem CA sequences adjacent to the MHV RNA poly(A) tail is most intriguing in this regard (Figs. 7C and S6). The CA sequences adjacent to the MHV RNA poly(A) tail are conserved in group 2 coronaviruses, present in both genomic and sg mRNAs, and positioned adjacent to the poly(A) template used for the initiation of negative-strand RNA synthesis (68). The coronavirus polymerase, nsp12, with nsp7 and 8 cofactors (75), initiates negative-strand RNA synthesis on the poly(A) tail of genomic RNA, leading to the synthesis of poly(U) at the 5′ end of negative-strand RNA. Because coronavirus nsp12 is a primer-dependent RNA polymerase (76), nsp8 is thought to prime negative-strand RNA synthesis (77), making a poly(U) product from the poly(A) tail of genomic RNA templates (16). Cleavage of the tandem CA sequences adjacent to the MHV RNA poly(A) tail would disrupt negative-strand RNA synthesis at the point of initiation. This provides a theoretically appealing mechanism for EndoU and other endoribonucleases to prevent the synthesis of dsRNA (Fig. 9).

MHV RNA was cleaved by one or more unspecified endoribonucleases in EndoU^mut^-infected RNase L^-/-^ BMM. Thus, in addition to EndoU- and RNase L-dependent cleavage of MHV RNA, we observed EndoU- and RNase L-independent cleavage of MHV RNA (Fig. 2 and Fig. S8). By using fold-change analyses between wt and mutant conditions, we attributed the majority of endoribonucleolytic cleavage sites in MHV RNA to EndoU activity and RNase L activity (Fig. 5); however, a substantial amount of cleavage in MHV RNA persisted in EndoU^mut^-infected RNase L^-/-^ BMM (Fig. 2, EndoU^mut^, red bars). More than 5% of the cyclic phosphates in EndoU^mut^-infected RNase L^-/-^ BMM RNA samples were in MHV RNA (Fig. 2, EndoU^mut^, red bars). This EndoU- and RNase L-independent cleavage of MHV RNA occurred predominantly at UA and CA dinucleotides (Fig. 4D); especially within IFNAR^-/-^ (Fig. S3A, Position −1 to +1) and RNase L^-/-^ (Fig. S3B, Position −1 to +1) BMM. Thus, the EndoU- and RNase L-independent cleavage of MHV RNA exhibited a nucleotide specificity similar to that of EndoU-dependent cleavage. It is possible that the H277A mutation in EndoU fails to completely inhibit endoribonuclease activity; however, we suspect that RNase A family members are responsible for this residual EndoU-independent cleavage of MHV RNA at U^⬇^A and C^⬇^A sequences. RNase A family enzymes are expressed in macrophage (63) and they cleave RNA at U^⬇^A and C^⬇^A sequences (62, 64). EndoU-independent cleavage of MHV RNA at the ^31332^C^⬇^AC^⬇^A^31335^ sequence was evident in WT BMM (Fig. S6A), IFNAR^-/-^ BMM (Fig. S6B) and RNase L^-/-^ BMM (Fig. S6C). The expression of RNases 4 and 5 (Fig. S12C) is consistent with residual cleavage at the ^31332^C^⬇^AC^⬇^A^31335^ sequences in EndoU^mut^-infected BMM (Fig. S6). These data indicate that the ^31332^C^⬇^AC^⬇^A^31335^ sequence in MHV RNA was susceptible to both EndoU-dependent and EndoU-independent cleavage. The atomic structure of EndoU revealed an RNase A-like catalytic domain (6); however, we did not anticipate the degree of overlap in substrate specificity observed for EndoU-dependent and EndoU-independent (presumably RNase A family) enzymes within BMM. Additional experiments will be required to address the identity and functional significance of the EndoU-independent (presumably RNase A family) enzymes within BMM.

### EndoU activity and cellular RNAs

Substantial amounts of EndoU-dependent and RNase L-dependent cleavage of MHV RNA were detected (Fig. 5), along with RNase L-dependent cleavage of rRNA (Figs. 8A, 8B and S10), but EndoU-dependent cleavage of cellular RNAs was not evident in our datasets. EndoU-dependent cleavage sites in rRNAs would be relatively easy to detect due to the abundance of rRNAs and to the well-established fold-change analyses proven to detect RNase L-dependent cleavage sites. Thus, we are confident that EndoU did not produce detectable cyclic phosphate moieties in rRNAs under the conditions of our experiments. Whether or not EndoU targets cellular mRNAs for cleavage is less certain. The low abundance of individual cellular mRNAs in our cyclic phosphate cDNA libraries precludes definitive assignment of one or another endoribonuclease to individual cleavage sites in individual cellular mRNAs. Thus, cellular mRNAs are cleaved by endoribonucleases, as they constitute ∼5% of all cleavage in our cyclic phosphate cDNA libraries (Figs. 2 and S8); however, we are not able to specify which endoribonucleases are responsible for individual cleavage sites within individual cellular mRNAs due to the limited abundance of any one cellular mRNA. Because EndoU-dependent cleavage sites were abundant in MHV RNAs, we suspect that EndoU is localized within RNA replication complexes, consistent with another report (21).

### Does nsp16 (2′ O-MT) regulate EndoU?

Deng and Baker highlight another unanswered questioned in the field (72): How is EndoU activity regulated to avoid unwanted cleavage events? This is an important question because MHV RNA integrity is critical for viral mRNA translation and viral RNA replication (Fig. 9). When EndoU cleaves MHV RNA, it must do so in a regulated manner to avoid self-destruction. Residual amounts of MHV genomic RNA must be maintained within infected cells to sustain an infection. One factor thought to regulate EndoU is nsp16, a 2′ O-methyltransferase (1).

When EndoU was first characterized, Ivanov and colleagues demonstrated that EndoU-mediated cleavage of RNA substrates was prevented by 2′-O-methylation (1). They also highlighted the modular nature of viral evolution, drawing attention to the side-by-side nature of nsp15 (EndoU) and nsp16 (2′ O-MT) within nidovirus genomes, suggesting a functional interplay between the two enzymes (1). 2′-O-methyltransferases have been functionally characterized in two families of positive-strand RNA viruses, coronaviruses (25, 78) and flaviviruses (79–82). One function of these enzymes is to methylate the adenosine of 5′ cap structures in viral mRNAs (81), to evade the antiviral activity of IFIT1 (25, 79, 80). Whether these enzymes can methylate other residues throughout viral RNA is less certain; however, 2′-O-methyltransferases are reported to inhibit the recognition of viral dsRNA by MDA5 (25). It is intriguing to note that EndoU cleavage sites (C^⬇^A and U^⬇^A sequences) contain adenosine. 2′-O-methylation of the pyrimidine at cleavage sites would prevent cleavage of viral RNA because the 2′ hydroxyl of ribose is the nucleophile responsible for attacking the phosphodiester backbone (59). Whether 2′ O methylation of adenosine can prevent EndoU-mediated cleavage of C^⬇^A and U^⬇^A sequences remains to be determined; however, some amount of intact MHV genomic RNA must be maintained within infected cells to sustain an infection.

RNAseq showed that MHV RNAs were abundant (Fig. S5) and there were proportional amounts of each MHV mRNA within infected cells (Fig. 6D) despite profound changes in endoribonuclease activity from one condition to another. Thus, neither EndoU nor RNase L activities were associated with extreme changes in the proportions of one MHV mRNA to another. Rather, relatively subtle changes in MHV mRNA1-8 proportions were observed. These data suggest that EndoU and RNase L activities modulate MHV RNA abundance during infections, but do not contribute to extreme changes in the relative amounts of one MHV mRNA to another. In contrast, the absence of EndoU activity during MHV infection lead to profound increases in host gene expression associated with response to dsRNA (Figs. 8D and 8E), despite the activation of RNase L activity. Expression and translation of cellular mRNAs occurs in the context of activated RNase L despite its’ ongoing degradation of cellular RNAs during a dsRNA-activated stress response (83, 84). Ongoing expression and translation of MHV mRNAs likely occur in the context of EndoU or RNase L activities in the same manner. When pre-existing host or viral mRNAs are destroyed by EndoU or RNase L activities, new MHV mRNAs are synthesized to refresh the pool of viral mRNAs. Thus, MHV replication can clearly tolerate - and perhaps benefit from - both EndoU and RNase L activities.

### Do EndoU and RNase L co-regulate MHV RNA gene expression and replication?

Importantly, EndoU and RNase L share a common cleavage site, UA (Fig. 4). Furthermore, we can distinguish between EndoU-dependent and RNase L-dependent cleavage of UA sequences because EndoU cleaves between U^⬇^A dinucleotides whereas RNase L cleaves after UA^⬇^ dinucleotides. Under some conditions, such as MHV^(S)^-infected and MHV^(V)^-infected WT BMM, UA sequences in viral RNA were cleaved predominantly by EndoU (Fig. 4G). Under other circumstances UA sequences in MHV RNA were cleaved predominantly by RNase L, as in PDE^mut^-infected and EndoU^mut^-infected WT BMM (Fig. 4G). In both cases, regardless of whether the host or viral endoribonuclease cleaves MHV RNA, the consequence will be an inhibition in viral mRNA translation and an inhibition in viral RNA replication (Fig. 9). It is interesting to see that both a host and a viral endoribonuclease have the capacity to inhibit magnitudes of MHV gene expression and replication by targeting a common set of UA sequences within the viral genome. It is also interesting that EndoU activity was subdued within IFNAR^-/-^ and RNase L^-/-^ cells, as if EndoU activity was modulated by RNase L activity (Fig. 5C). Together, these results suggest an interesting interplay between EndoU and dsRNA-activated host responses (Fig. 9).

### Summary

We addressed a key question in the field (72): What is the natural target of coronavirus EndoU? We find that EndoU targets MHV RNA within infected cells, cleaving viral RNA on the 3′ side of pyrimidines with a strong preference for cleavage between U^⬇^A and C^⬇^A sequences (endoY^⬇^A). We postulate that EndoU cleaves MHV RNA in a regulated manner, to inhibit negative-strand RNA synthesis, reducing the accumulation of viral dsRNA, while ensuring continuing virus replication (Fig. 9). By regulating the synthesis and accumulation of viral dsRNA, coronaviruses can evade double-stranded RNA-activated antiviral responses within infected cells (22, 26, 27, 72).

## ACKNOWLEDGMENTS

We thank members of the Barton and Weiss labs for critical evaluation of the manuscript. We thank Evan Lester for assistance with bioinformatics analyses. This work was supported by Public Health Service grants from the National Institutes of Health (F30 AI140615 to RA, AI104887 to SRW, GM119550 to JRH and AI042189 to DJB) and the Swiss National Science Foundation (SNF project 149784).

## AUTHOR CONTRIBUTIONS

Rachel Ancar: Experimental Design, Experimental Procedures, Bioinformatics, Data Analysis, Data Curation, Interpretation of Data and Manuscript Preparation.

Yize Li: Experimental Design, Experimental Procedures and Interpretation of Data.

Eveline Kinder: Experimental Design and Experimental Procedures.

Daphne Cooper: Methodology and Pilot Study.

Monica Ransom: Experimental Procedures.

Volker Thiel: Experimental Design, Project Administration, Funding Acquisition and Data Interpretation.

Susan Weiss: Experimental Design, Project Administration, Funding Acquisition and Data Interpretation.

Jay Hesselberth: Experimental Design, Project Administration, Funding Acquisition and Data Interpretation and Manuscript Preparation.

David Barton: Experimental Design, Project Administration, Funding Acquisition, Data Interpretation and Manuscript Preparation.

## Competing Interests

Authors report no competing interests.

## SUPPLEMENTAL FIGURES AND TABLES

**Figure S1.** Frequency and location of endoribonuclease cleavage sites in U6 snRNA and MHV antigenomic RNA in WT, IFNAR^-/-^, and RNase L^-/-^ BMM. Normalized 2’-3’-cp cDNA reads captured at each position along the (A) U6 snRNA and (B) MHV antigenomic RNA at 9 and 12 hpi with MHV^(S)^, MHV^(V)^, PDE^mut^, and EndoU^mut^ virus in IFNAR^-/-^ BMM.

**Figure S2. Dinucleotide cleavage pattern in MHV genomic RNA downstream of captured 3’ RNA end.** Percent total 2’-3’-cp cDNA reads captured at each dinucleotide in WT BMM at 9 and 12 hpi for bases +1:+2 and +2:+3 from the captured cleavage position (0-base).

**Figure S3. Endoribonuclease cleavage preferences in MHV RNA from IFNAR^-/-^ and RNase L^-/-^ BMM. (**A) and (B) Dinucleotide specificity analysis for cleavage in MHV genomic RNA by percent total cyclic phosphate cDNA reads captured at each 3’-dinucleotide at 9 and 12 hpi in (A) IFNAR^-/-^ BMM for positions −2:-1 and in (B) RNase L^-/-^ BMM for positions −1:+1.

**Figure S4. Interaction between RNase L and EndoU cleavage at UA sequences in MHV RNA.** (A) UA cleavage scoring analysis. All UA sequences in the MHV genomic RNA with >= 30 cyclic phosphate counts in either the UA^⬇^ or U^⬇^A cleavage position were compared by calculating the ratio of normalized counts (UA^⬇^ counts /U^⬇^A counts). Ratios > 1 were scored as UA^⬇^ (RNase L) sites and ratios <1 were scored as U^⬇^A sites (EndoU) and total number of scored sites for either position are shown for each condition of viral infection in IFNAR^-/-^ and RNase L^-/-^ BMM. (B) Frequency and location of UA positions in WT BMM under all conditions of viral infection which had ≥ 50 counts at U^⬇^A positions and ≤ 1 counts at UA^⬇^ positions. The top 5 of these positions by normalized count are labeled. (C) Frequency and location of UA positions in IFNAR^-/-^ and RNase L^-/-^ BMM under all conditions of viral infection which had ≥ 50 counts at U^⬇^A positions and ≤ 1 counts at UA^⬇^ positions.

**Figure S5. MHV RNA abundance and RNAseq reads across the MHV genome.** (A) Total RNAseq normalized counts assigned to MHV genomic features for each library. (B) Coverage of RNAseq read density across the MHV genome in reads per million.

**Figure S6. Frequency and location of endoribonuclease cleavage in MHV 3’-UTR.** Nucleotide and sequence resolution graphs displaying normalized counts in (A) WT, (B) IFNAR^-/-^ (C) RNase L^-/-^ BMM during infection with MHV^(V)^ and EndoU^mut^ MHV. Nucleotide resolution graphs of region directly upstream of poly-A tail with cyclic phosphate counts normalized by RNA abundance in (D) WT, (E) IFNAR^-/-^ (F) RNase L^-/-^ BMM.

**Figure S7. Regional cleavage of MHV RNA and total subgenomic mRNA abundance.** (A) Sum of cyclic phosphate counts normalized by mRNA abundance displayed by region for WT, IFNAR^-/-^, RNase L^-/-^ BMM across all conditions of viral infection at 12 hpi. (B) Percent of sum of normalized counts per length of genomic region for WT, IFNAR^-/-^, RNase L^-/-^ BMM across all conditions of viral infection at 12 hpi. Dotted line represents baseline percent of cleavage expected by cell type ([total number of cyclic phosphate counts / total genome size x 100]). (C) Sum of all subgenomic mRNAs (RPM) for WT, IFNAR^-/-^, RNase L^-/-^ BMM across all conditions of viral infection at 9 and 12 hpi.

**Figure S8. Cyclic phosphate sequencing analysis of experiment 2.** (A) Normalized cyclic phosphate cDNA reads ([reads at each position / total reads in library]) aligning to host and viral RNAs at 9 and 12 hpi in WT and RNase L^-/-^ bone marrow macrophages (BMM). (B) Putative cleavage sites attributed to EndoU or RNase L were calculated from RNase L- or EndoU-dependent signal generated by subtracting signal from each captured position that occurs in the absence of either enzyme (RNase L^-/-^ BMM or during EndoU^mut^ infection). These data were then filtered for sites with reads representing at least 0.01 % of total reads in the library. At each of these positions, the log2 fold change in signal when either RNase L or EndoU were absent was calculated and sites with >= 2.5 fold change were called as putative RNase L or EndoU sites. (C and D) Dinucleotide specificity analysis for cleavage in MHV genomic RNA by percent total cDNA reads captured at each 3’-dinucleotide in WT BMM at 9 and 12 hpi for (C) Dinucleotide analysis for positions −2: −1 and (D) Dinucleotide analysis for positions −1:+1 from captured cleavage position (0 position). (E) Cumulative distribution of normalized counts by position of the MHV genome for every position with >= 10 cyclic phosphates counts across all cell types and infection conditions. (F) UA cleavage scoring analysis. All UA sequences in the MHV genomic RNA with ≥ 30 cyclic phosphate counts in either the UA^⬇^ or U^⬇^A cleavage position were compared by calculating the ratio of normalized counts (UA^⬇^ counts /U^⬇^A counts). Ratios > 1 were scored as UA^⬇^ (RNase L) sites and ratios <1 were scored as U^⬇^A sites (EndoU) and total number of scored sites for either position are shown for each condition of viral infection in WT BMM at 9 and 12 hpi. (H) Model of EndoU and RNase L interaction at UA sites in MHV RNA. (G) Nucleotide and sequence resolution graphs displaying normalized counts in WT BMM during infection with MHV^(S)^ and EndoU^mut^ MHV of region directly upstream of poly(A) tail.

**Figure S9. Cyclic phosphate and RNAseq analysis of MHV RNA from experiment 2.** (A) Sum of normalized counts displayed by region for WT and RNase L^-/-^ BMM across all conditions of viral infection at 12 hpi. Transcription regulatory sequences (TRS) are numbered by their associated mRNA (2–7). Other genomic regions are labeled as shown in Figure 1A. (B) Sum of cyclic phosphate counts normalized by mRNA abundance and length of each genomic region [(sum per region (cyclic phosphate counts / RNAseq counts) / length of region (bp) *100] sum of cyclic phosphate abundance normalized counts per region)/length of region * 100] displayed by region for WT and RNase L^-/-^ BMM across all conditions of viral infection at 12 hpi. (C) Normalized counts (sum of subgenomic mRNA / sum of all mRNAs) of subgenomic mRNAs detected in WT and RNase L^-/-^ BMM across all conditions of viral infection at 9 and 12 hpi. (D) Sum of all subgenomic mRNAs (RPM) for WT and RNase L^-/-^ BMM across all conditions of viral infection at 9 and 12 hpi.

**Figure S10. RNase L targeting of rRNA and mRNA during WT and mutant MHV infection.** (A) Top 3 RNase L-dependent cleavage sites in 18S rRNA by fold change (log2WT BMM / RNase L^-/-^ BMM) for all conditions of infection at 9 and 12 hpi in WT BMM. (B) Table of total RNase L- or EndoU-dependent cleavage sites in 18S, 28S, 5.8S, and 5S rRNA. RNase L- or EndoU-dependent cleavage were determined by identifying the top 1% of enzyme-dependent signal with a > 4 fold change in signal in the absence of RNase L or EndoU that match the RNase L (“UA”, “UU”, “UG”, “UC”) or EndoU (“UA”, “CA”) sequence preferences. (C) Distribution of sites in rRNA by fold change in signal when comparing WT or IFNAR^-/-^ BMM mock-infected or virus-infected with MHV^(S)^, MHV^(V)^, PDE^mut^, and EndoU^mut^ virus to RNase L^-/-^ BMM. Log2 fold change in the absence of RNase L activity was calculated for each position in the rRNA. The distribution of the top 100 RNase L-dependent sites were compared in WT and IFNAR^-/-^ BMM under conditions of mock infection or virus infection with MHV^(S)^, MHV^(V)^, PDE^mut^, and EndoU^mut^ virus at 12 hpi. (D) Distribution of sites in 18S, 28S, 5S, and 5.8S rRNA with the greatest fold change in signal when comparing mock infection or virus infection with MHV^(S)^, MHV^(V)^, PDE^mut^ virus to infection with EndoU^mut^ virus. Log2 fold change in the absence of EndoU activity was calculated for each position in the rRNA. The distribution of the top 100 EndoU-dependent sites were compared in all cell types across conditions of mock infection or infection with MHV^(S)^, MHV^(V)^, PDE^mut^, and EndoU^mut^ virus at 9 and 12 hpi.

**Figure S11. Correlation between cyclic phosphate counts and RNA abundance.** (A and B). Correlation between mRNA abundance and 2’-3’-cp counts at each base captured in the MHV genome (all comparisons significant (p < 10^^50^) for experiment 1 (A) and experiment 2 (B). (C and D) Distribution of sites in MHV by normalized cyclic phosphates counts for sense and antisense RNAs from experiment 1 (C) and experiment 2 (D) across all conditions of infection and cell types. Positions are only shown if there was > 1 read in either the sense and antisense RNA.

**Figure S12. Effect of WT and mutant MHV infection on other cellular endoribonucleases.** (A and B) Volcano plot of differentially expressed (FDR < 0.05) and upregulated (logFC > 2)/ downregulated (logFC < −2) genes when comparing EndoU^mut^ infection to mock infection in WT BMM (A) or MHV^(s)^ infection to EndoU^mut^ infection in WT BMM. (C) Expression (log10 normalized RNAseq counts) of RNase A, angiogenin, and RNaseT2 genes with at least 5 > normalized counts in all conditions.

**Table S1 and S2. Dinucleotide enrichment and de-enrichment analysis at 9 and 12 hpi in WT BMM.** Complete tables of dinucleotide enrichment and de-enrichment for −2 base: +2 base (table 1) or −1 base:-1 base (table 2) from the captured RNA end (0-base position) for each condition of viral infection at 9 and 12 hpi in WT BMM by comparing the frequency of dinucleotide capture in experimental conditions to the frequency of occurrence for each dinucleotide in the MHV genomic RNA sequence (control). Significant enrichment was determined by adjusted p-value (q) for fold change (log2(experiment / control)). (<0.02*, <0.0001**, <1×10^8***^).

**Table S3: SNP variants in MHV genome related to endoribonuclease cleavage.** Table of all single nucleotide variants (SNPs) identify from alignments of RNAseq libraries to the MHV genome. The SNPs in **green** are sites where the mutation generated a “CA” dinucleotide that was cleaved by EndoU. The SNPs in **red** and **yellow** are the inactivating mutations in the EndoU and PDE domains of MHV respectively.

## Notes

### Competing Interest Statement

The authors have declared no competing interest.

