## Supplemental Figures for "Physiologic RNA Targets and Refined Sequence Specificity of Coronavirus EndoU"

A

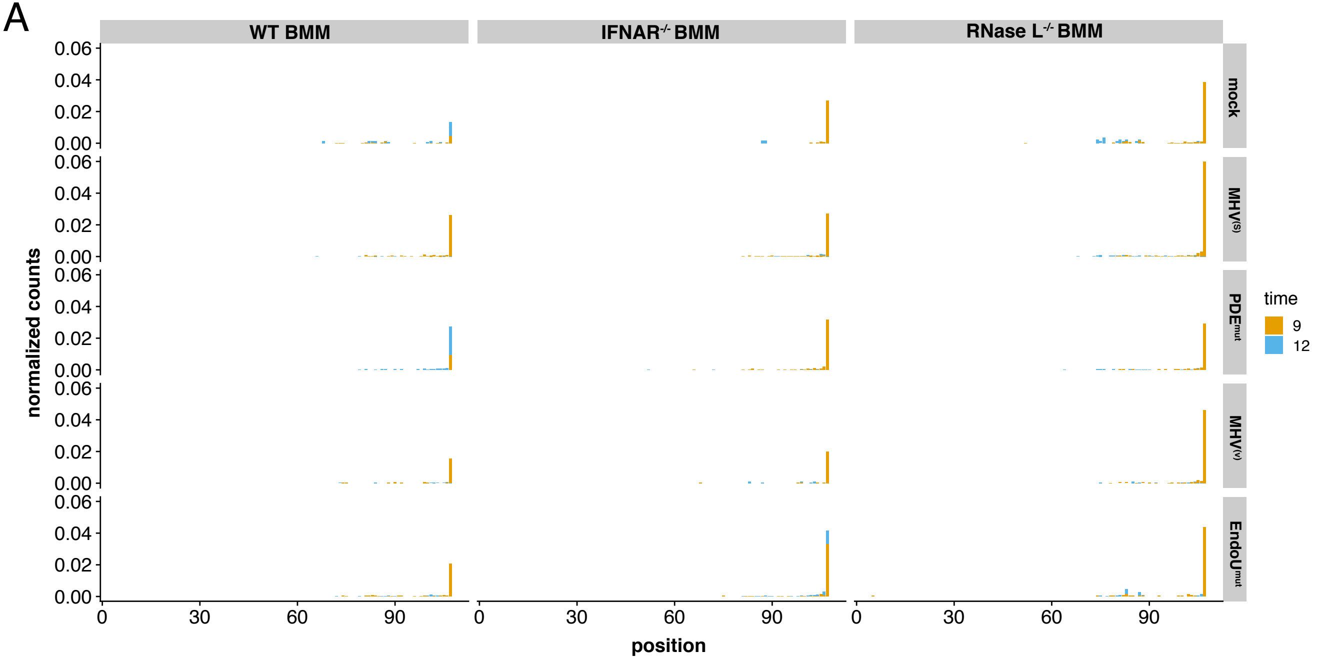
WT BMM
IFNAR-/- BMM
RNase L-/- BMM
mock
MHV(S)
PDEmut
MHV(V)
EndoUmut
normalized counts
0.06
0.04
0.02
0.00
0
30
60
90
position
time
9
12

B

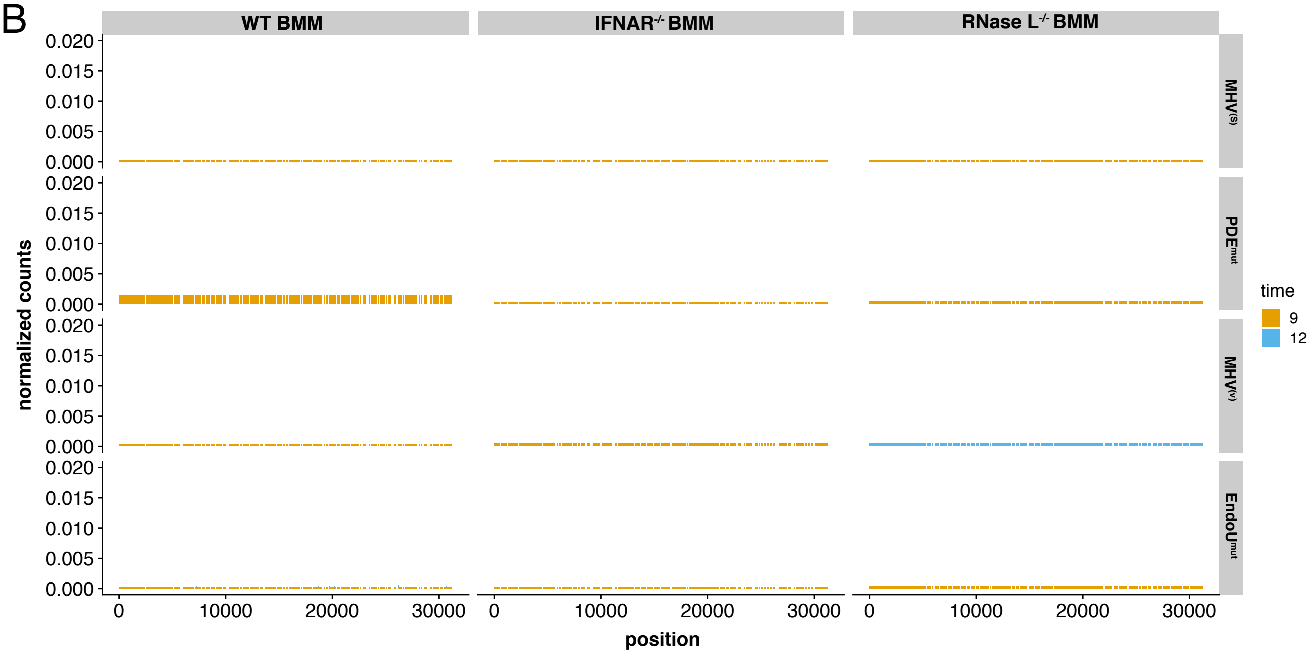
WT BMM
IFNAR-/- BMM
RNase L-/- BMM
MHV(S)
PDEmut
MHV(V)
EndoUmut
normalized counts
0.020
0.015
0.010
0.005
0.000
0
10000
20000
30000
position
time
9
12

Figure S1

A **Position +1 to +2 dinucleotide analysis in WT BMM**

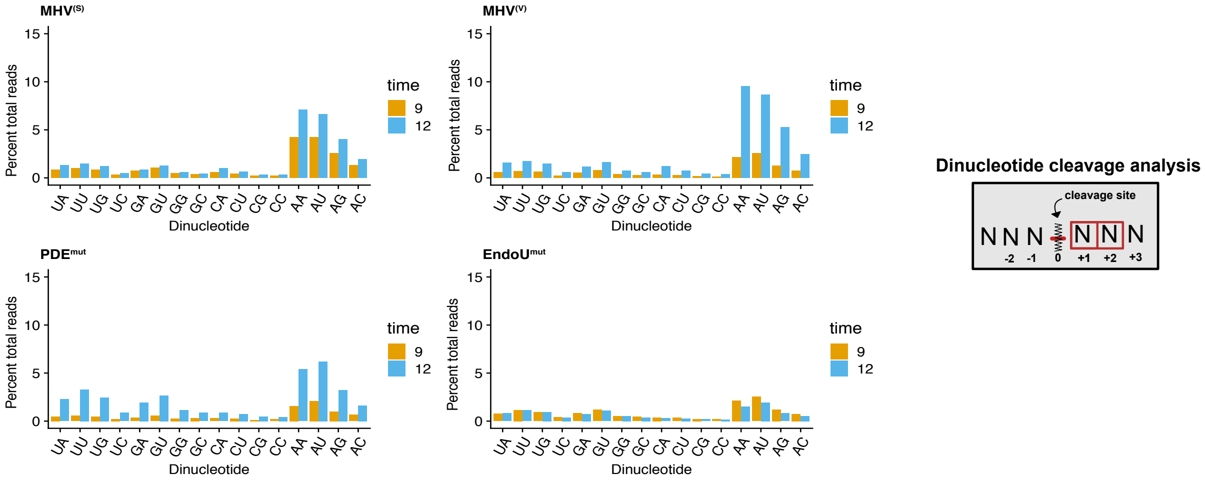


B **Position +2 to +3 dinucleotide analysis in WT BMM**

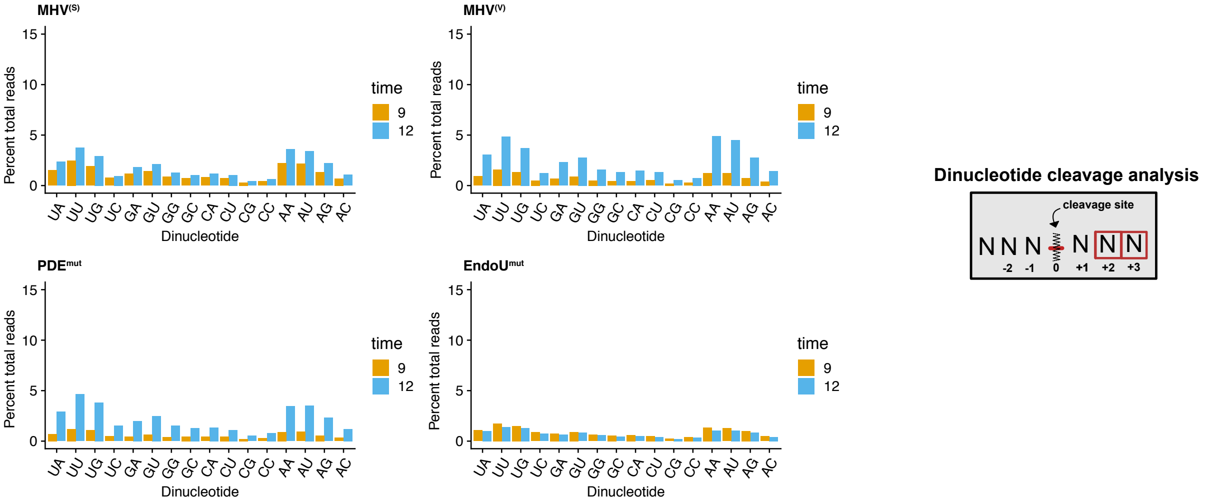


Figure S2

#### A Dinucleotide analysis in IFNAR$^{-/-}$ BMM

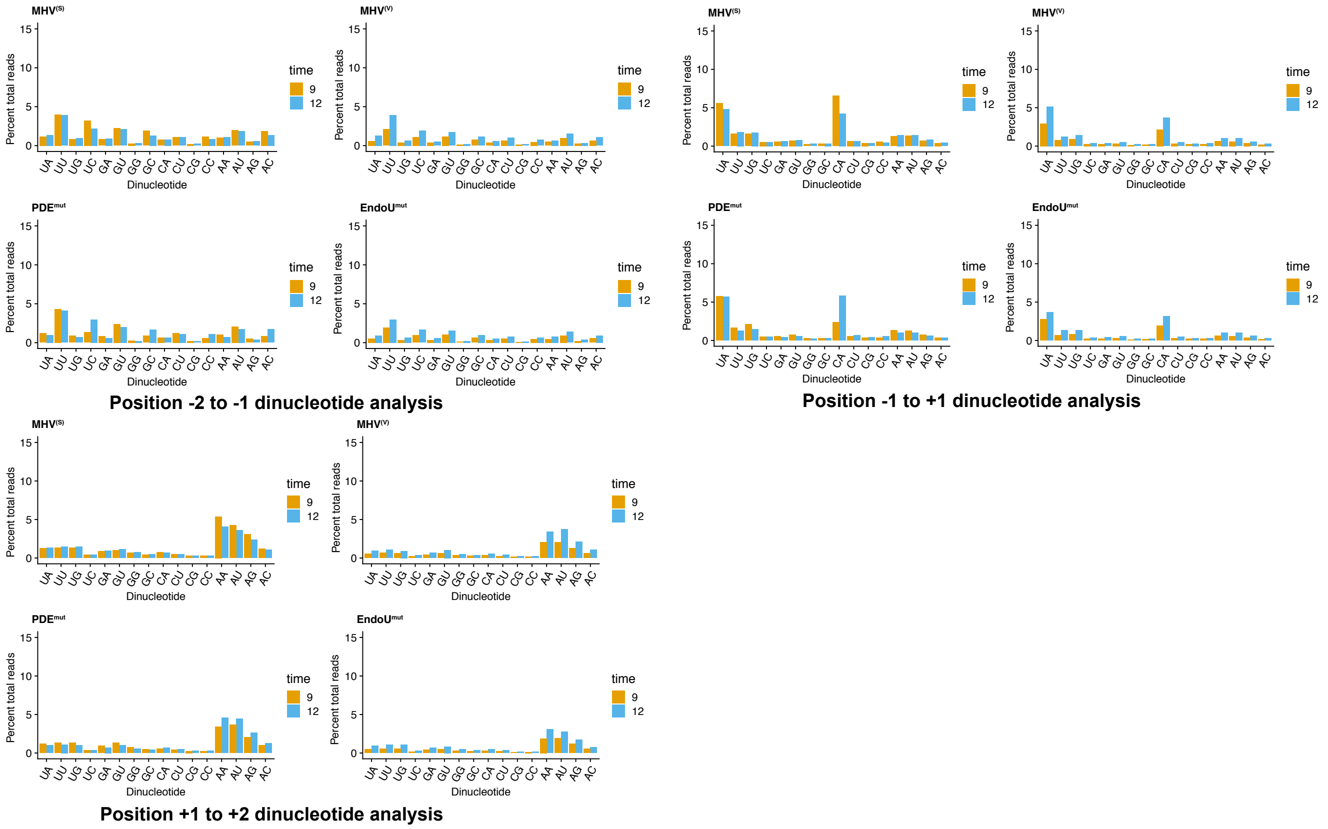


#### B Dinucleotide analysis in RNase L$^{-/-}$ BMM

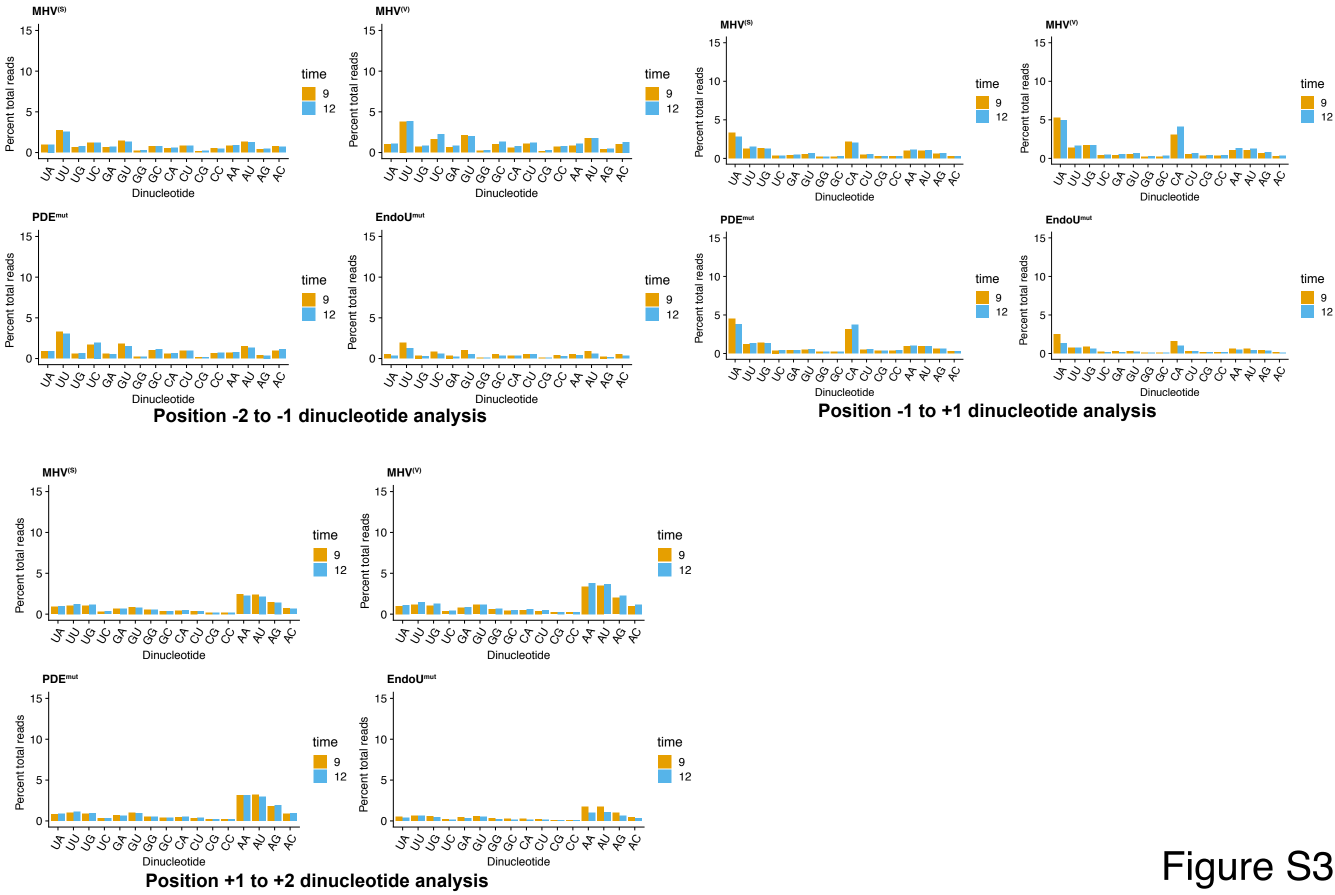


Figure S3

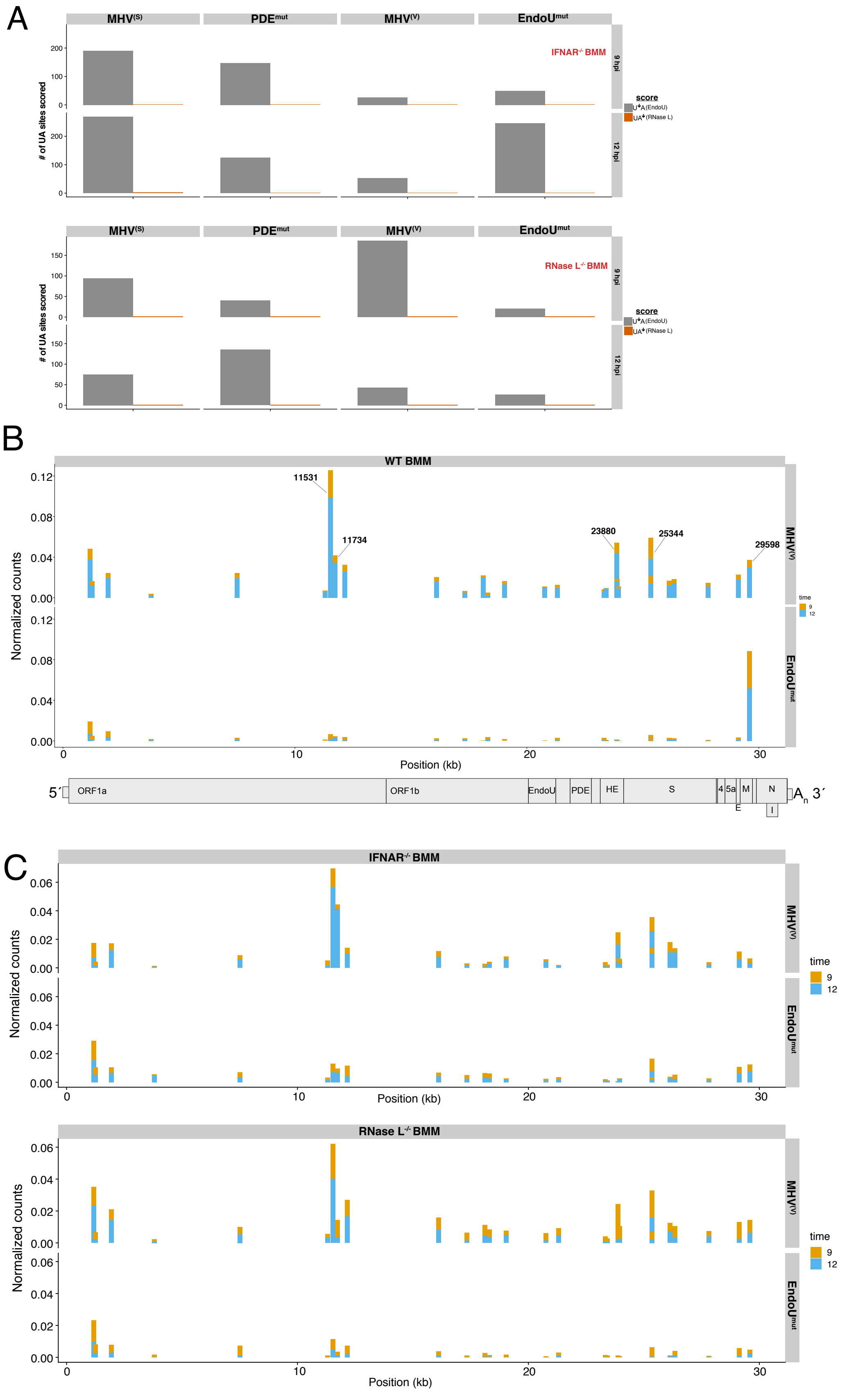
A
MHV(S)
PDEmut
MHV(V)
EndoUmut
IFNAR-/- BMM
RNase L-/- BMM
### of UA sites scored
9 hpi
12 hpi
score
U*A(EndoU)
UA*(RNase L)
B
WT BMM
11531
11734
23880
25344
29598
Normalized counts
Position (kb)
time
5´
ORF1a
ORF1b
EndoU
PDE
HE
S
4
5a
M
N
E
I
An 3´
C
IFNAR-/- BMM
RNase L-/- BMM

Figure S4

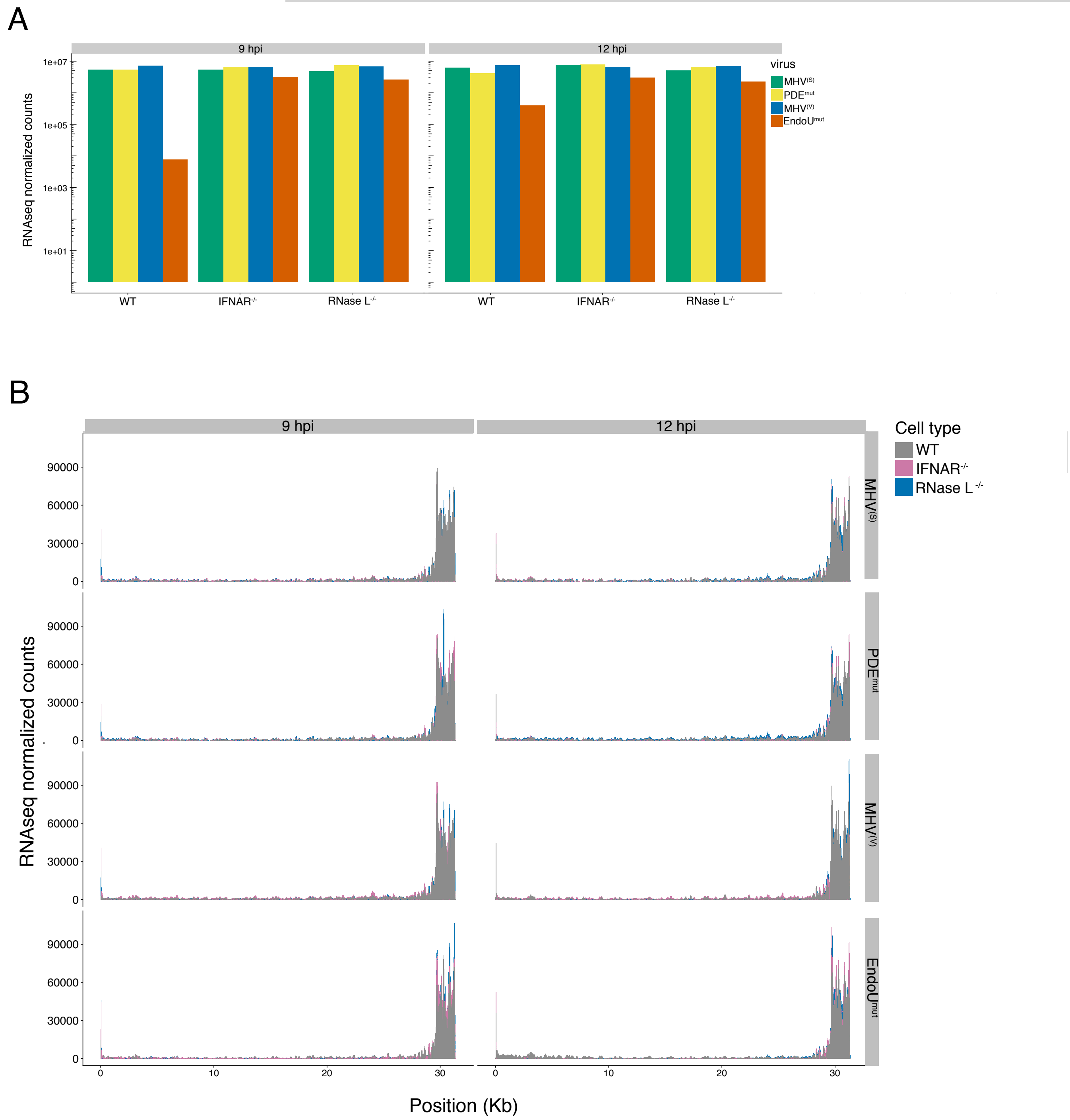
A
9 hpi
12 hpi
RNAseq normalized counts
1e+07
1e+05
1e+03
1e+01
WT
IFNAR-/-
RNase L-/-
virus
MHV(S)
PDEmut
MHV(V)
EndoUmut
B
Cell type
WT
IFNAR-/-
RNase L-/-
90000
60000
30000
0
MHV(S)
PDEmut
MHV(V)
EndoUmut
10
20
30
Position (Kb)

Figure S5

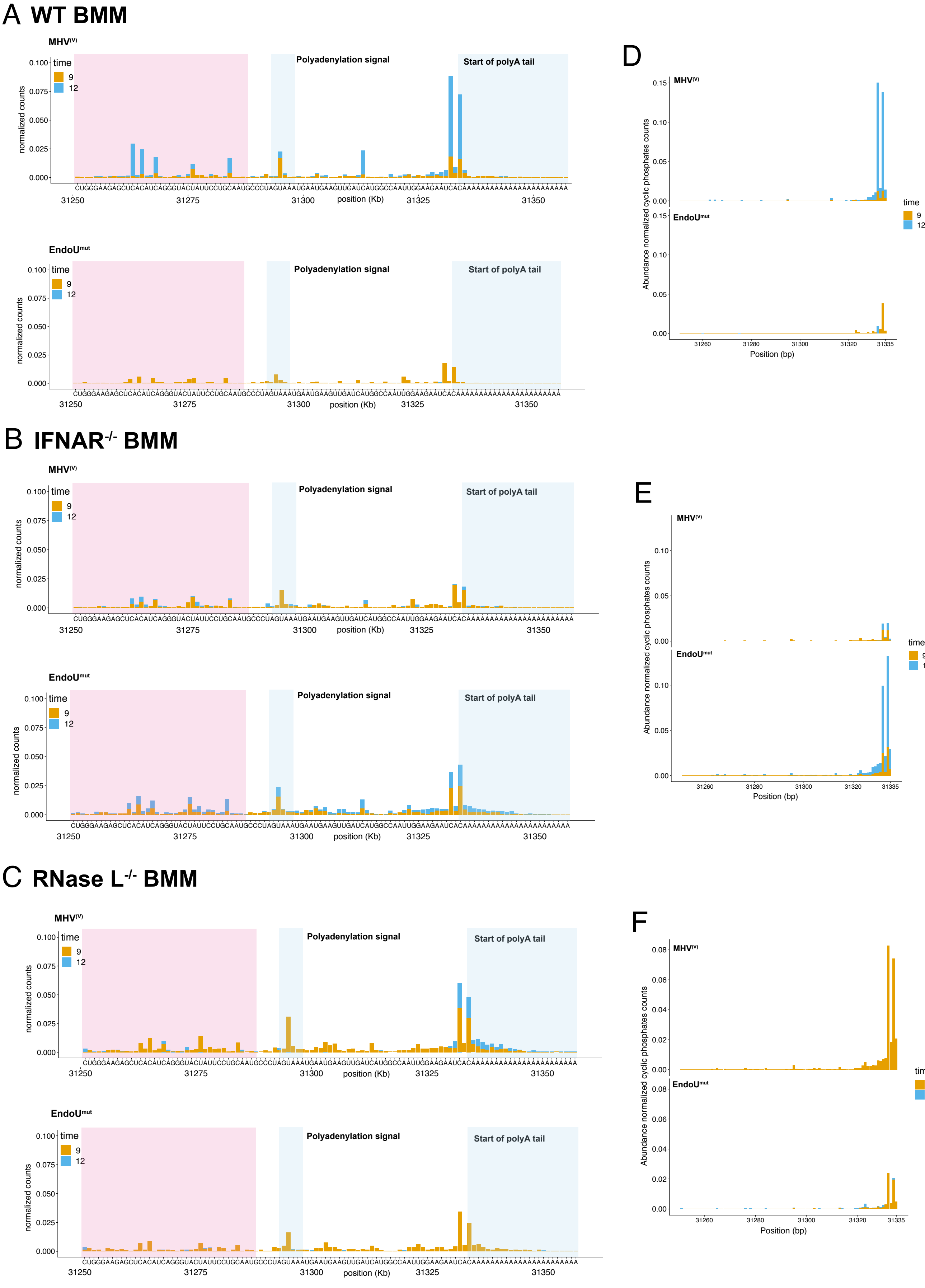
A WT BMM
MHV(V)
EndoUmut
B IFNAR-/- BMM
C RNase L-/- BMM
D
E
F
time
9
12
normalized counts
Polyadenylation signal
Start of polyA tail
CUGGGAAGAGCUCACAUCAGGGUACUAUUCCUGCAAUGCCCUAGUAAAUGAAUGAAGUUGAUCAUGGCCAAUUGGAAGAAUCACAAAAAAAAAAAAAAAAAAAAAAA
31250
31275
31300
31325
31350
position (Kb)
Abundance normalized cyclic phosphates counts
Position (bp)
31260
31280
31300
31320
31335

Figure S6

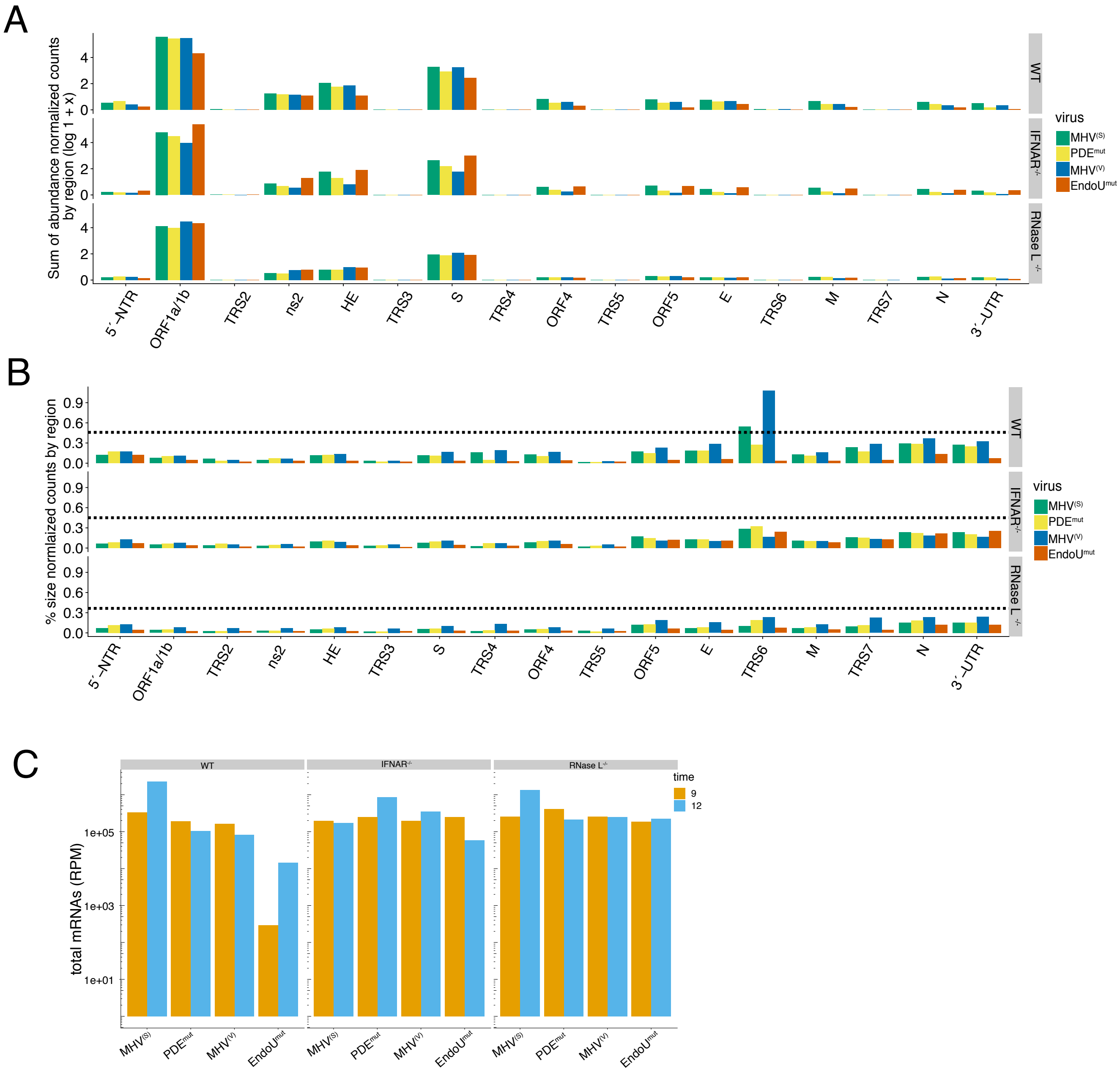
A
Sum of abundance normalized counts by region (log 1 + x)
WT
IFNAR-/-
RNase L -/-
virus
MHV(S)
PDEmut
MHV(V)
EndoUmut
5´-NTR
ORF1a/1b
TRS2
ns2
HE
TRS3
S
TRS4
ORF4
TRS5
ORF5
E
TRS6
M
TRS7
N
3´-UTR
B
% size normlaized counts by region
C
total mRNAs (RPM)
time
9
12

Figure S7

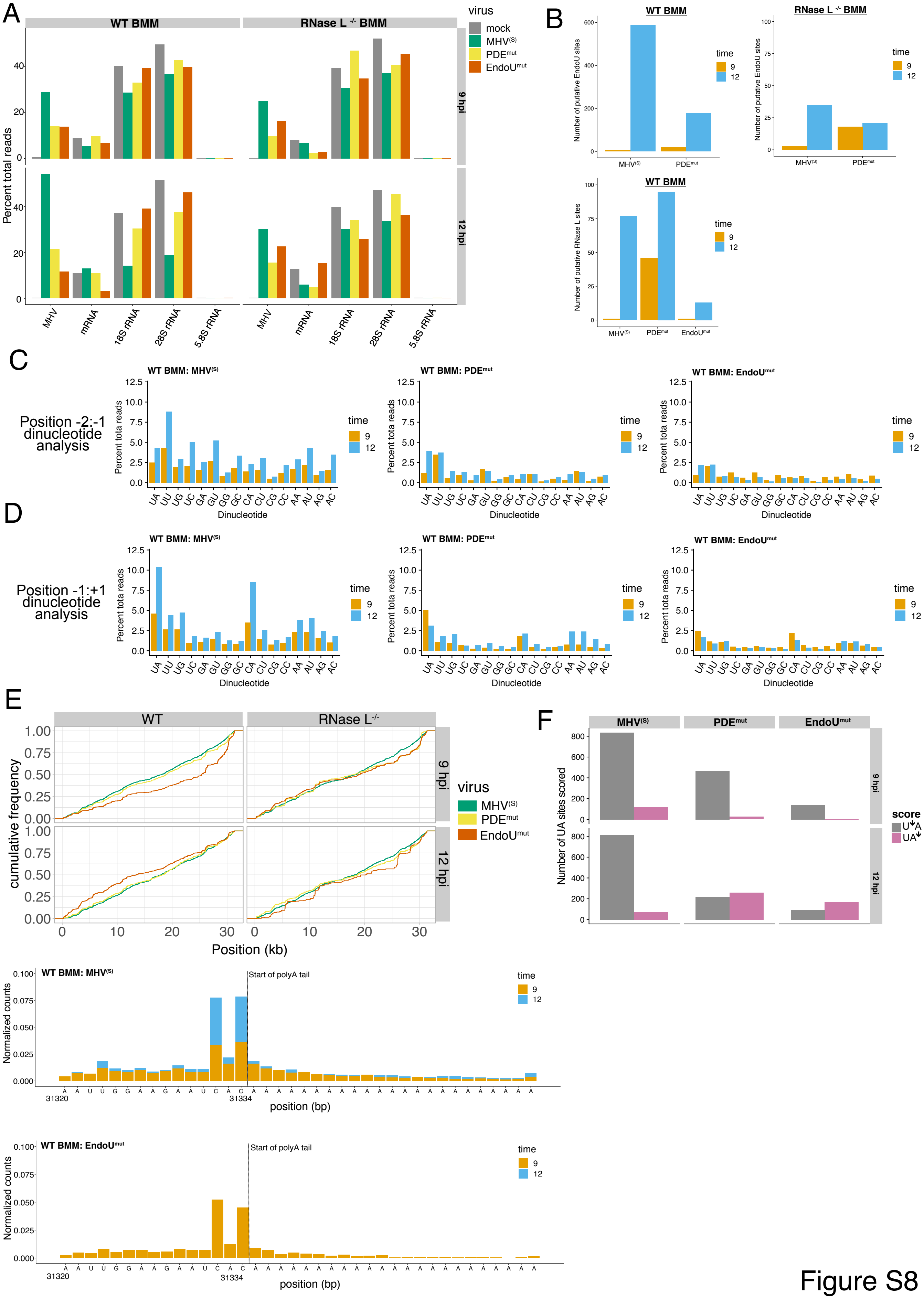
A
WT BMM
RNase L -/- BMM
virus
mock
MHV(S)
PDEmut
EndoUmut
9 hpi
12 hpi
Percent total reads
MHV
mRNA
18S rRNA
28S rRNA
5.8S rRNA
B
WT BMM
RNase L -/- BMM
Number of putative EndoU sites
Number of putative RNase L sites
time
9
12
MHV(S)
PDEmut
EndoUmut
C
Position -2:-1 dinucleotide analysis
WT BMM: MHV(S)
WT BMM: PDEmut
WT BMM: EndoUmut
Percent tota reads
Dinucleotide
UA UU UG UC GA GU GG GC CA CU CG CC AA AU AG AC
D
Position -1:+1 dinucleotide analysis
E
WT
RNase L-/-
cumulative frequency
Position (kb)
F
Number of UA sites scored
score
U↓A
UA↓
WT BMM: MHV(S)
Start of polyA tail
Normalized counts
position (bp)
A A U U G G A A G A A U C A C A A A A A A A A A A A A A A A A A A A A A A A
31320
31334
WT BMM: EndoUmut

Figure S8

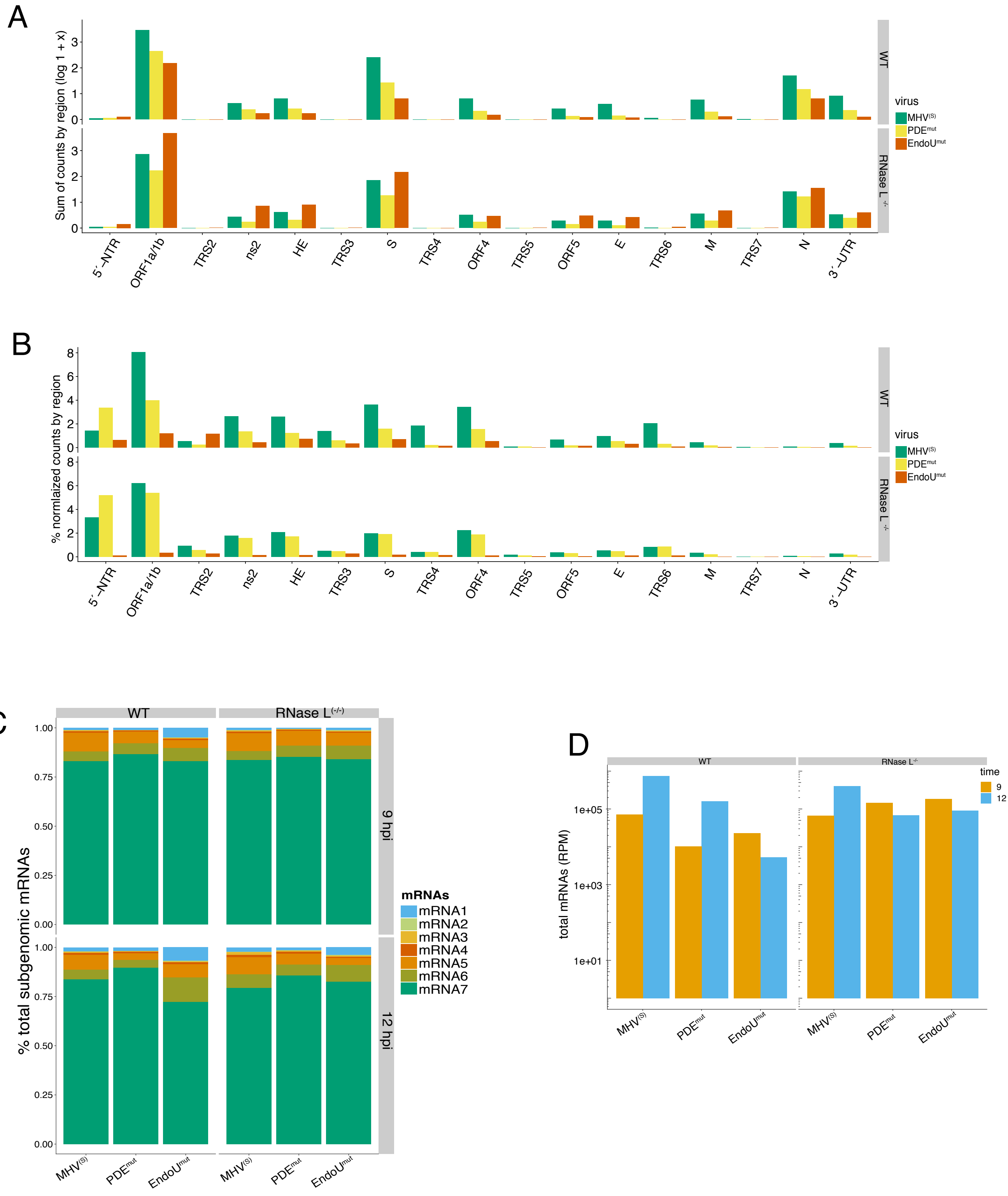
A
Sum of counts by region (log 1 + x)
WT
RNase L -/-
virus
MHV(S)
PDEmut
EndoUmut
5´-NTR
ORF1a/1b
TRS2
ns2
HE
TRS3
S
TRS4
ORF4
TRS5
ORF5
E
TRS6
M
TRS7
N
3´-UTR
B
% normlaized counts by region
C
WT
RNase L(-/-)
% total subgenomic mRNAs
9 hpi
12 hpi
mRNAs
mRNA1
mRNA2
mRNA3
mRNA4
mRNA5
mRNA6
mRNA7
MHV(S)
PDEmut
EndoUmut
D
total mRNAs (RPM)
time
9
12
1e+05
1e+03
1e+01

Figure S9

A

| Position | time | mock | $MHV^{(S)}$ | $MHV^{(V)}$ | $PDE^{mt}$ | $EndoU^{mt}$ |
|---|---|---|---|---|---|---|
| $UA^{772}$ | 9 | -1.09 | 3.47 | 4.48 | 6.58 | 8.66 |
| $UU^{771}$ | 9 | -0.65 | 1.32 | 2.65 | 3.77 | 7.30 |
| $UU^{543}$ | 9 | -2.05 | -0.07 | 1.00 | 1.94 | 4.29 |
| $UA^{772}$ | 12 | 2.21 | 3.29 | 1.02 | 5.52 | 4.88 |
| $UU^{542}$ | 12 | 4.32 | 2.01 | 2.17 | 3.37 | 4.26 |
| $UU^{771}$ | 12 | -0.64 | -0.33 | 1.55 | 1.57 | 3.94 |

B

| | RNase L sites | EndoU sites |
|---|---|---|
| 18S rRNA | 8 | 0 |
| 28S rRNA | 26 | 0 |
| 5.8S rRNA | 3 | 0 |
| 5S rRNA | 1 | 0 |

C

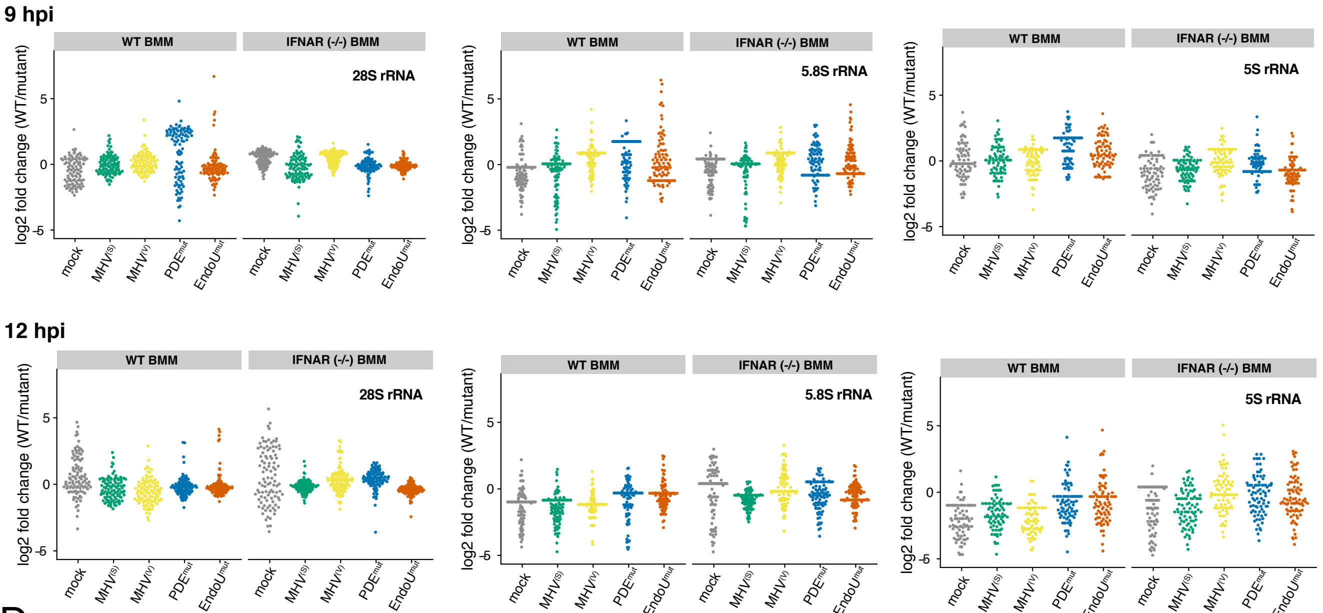


D

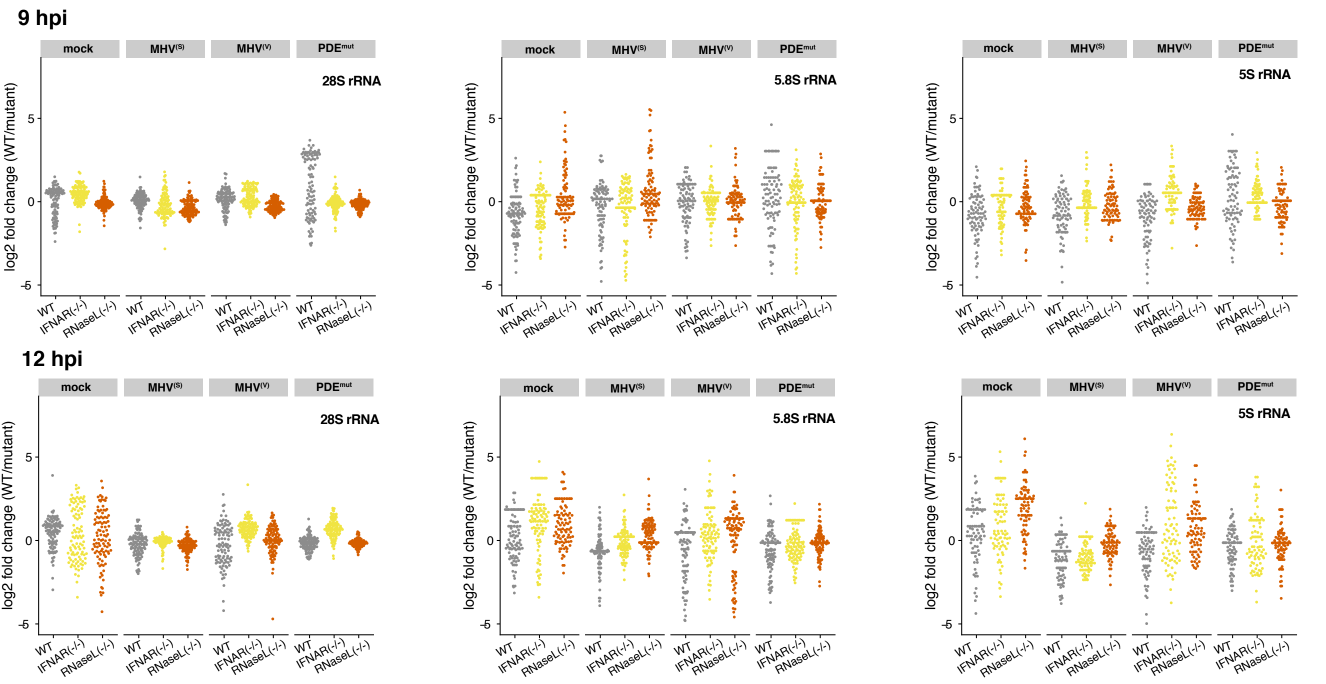


Figure S10

A

| Cell | Virus | Corr. estimate |
|---|---|---|
| WT | MHV$^{(S)}$ | 0.151 |
| WT | MHV$^{(V)}$ | 0.109 |
| WT | PDE$^{mut}$ | 0.084 |
| WT | EndoU$^{mut}$ | 0.139 |
| IFNAR $^{-/-}$ | MHV$^{(S)}$ | 0.194 |
| IFNAR $^{-/-}$ | MHV$^{(V)}$ | 0.103 |
| IFNAR $^{-/-}$ | PDE$^{mut}$ | 0.154 |
| IFNAR $^{-/-}$ | EndoU$^{mut}$ | 0.252 |
| RNaseL $^{-/-}$ | MHV$^{(S)}$ | 0.210 |
| RNaseL $^{-/-}$ | MHV$^{(V)}$ | 0.172 |
| RNaseL $^{-/-}$ | PDE$^{mut}$ | 0.174 |
| RNaseL $^{-/-}$ | EndoU$^{mut}$ | 0.229 |

B

| Cell | Virus | Corr. estimate |
|---|---|---|
| WT | MHV$^{(S)}$ | 0.151 |
| WT | PDE$^{mut}$ | 0.084 |
| WT | EndoU$^{mut}$ | 0.139 |
| RNaseL $^{-/-}$ | MHV$^{(S)}$ | 0.194 |
| RNaseL $^{-/-}$ | PDE$^{mut}$ | 0.103 |
| RNaseL $^{-/-}$ | EndoU$^{mut}$ | 0.154 |

C

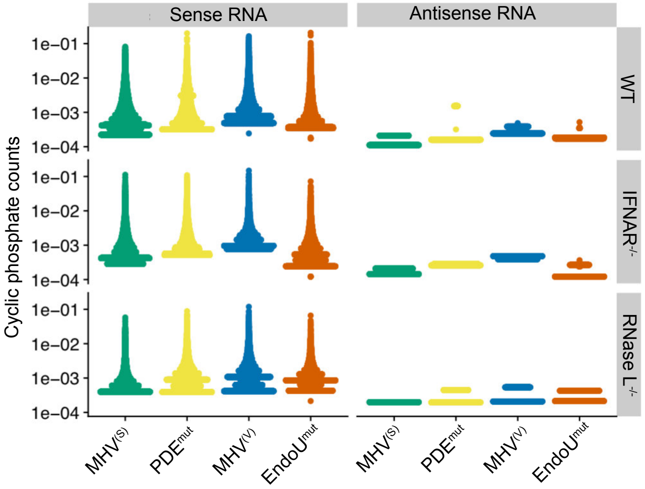


D

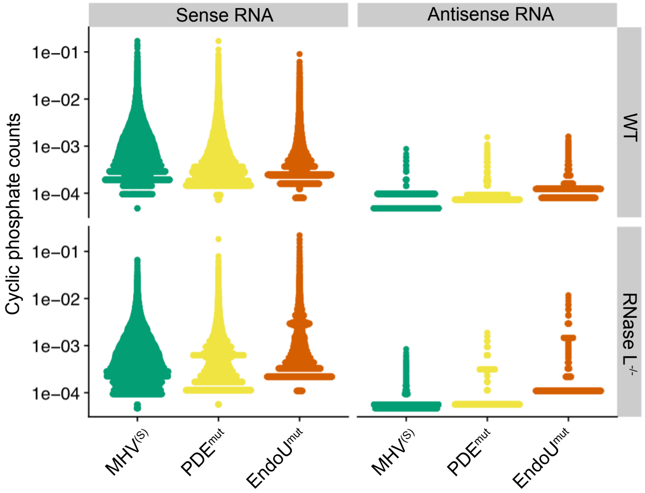


Figure S11

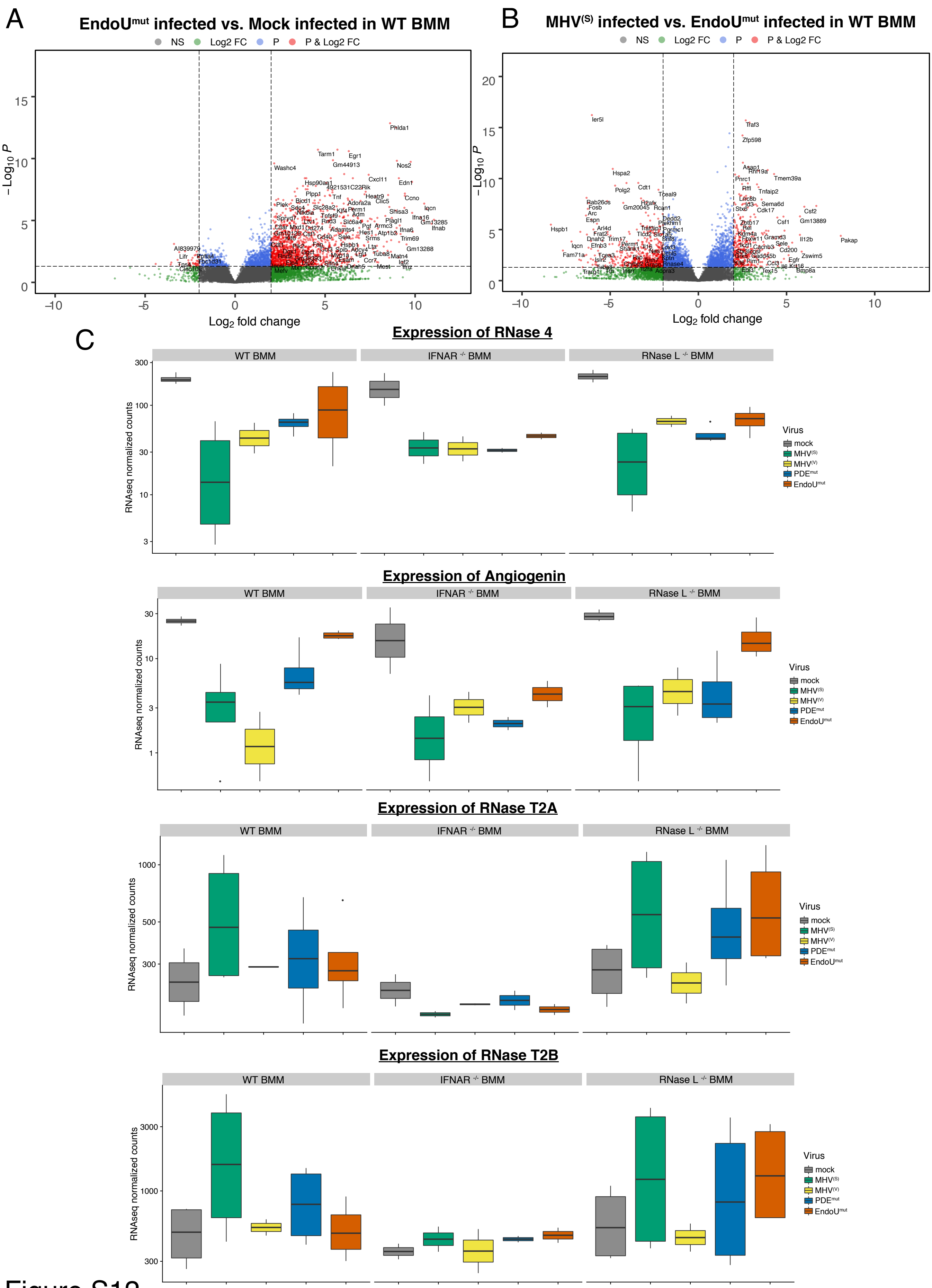
A
EndoU^mut infected vs. Mock infected in WT BMM
NS
Log2 FC
P
P & Log2 FC
−Log10 P
Log2 fold change
B
MHV(S) infected vs. EndoU^mut infected in WT BMM
NS
Log2 FC
P
P & Log2 FC
−Log10 P
Log2 fold change
C
Expression of RNase 4
WT BMM
IFNAR -/- BMM
RNase L -/- BMM
RNAseq normalized counts
Virus
mock
MHV(S)
MHV(V)
PDEmut
EndoUmut
Expression of Angiogenin
WT BMM
IFNAR -/- BMM
RNase L -/- BMM
RNAseq normalized counts
Virus
mock
MHV(S)
MHV(V)
PDEmut
EndoUmut
Expression of RNase T2A
WT BMM
IFNAR -/- BMM
RNase L -/- BMM
RNAseq normalized counts
Virus
mock
MHV(S)
MHV(V)
PDEmut
EndoUmut
Expression of RNase T2B
WT BMM
IFNAR -/- BMM
RNase L -/- BMM
RNAseq normalized counts
Virus
mock
MHV(S)
MHV(V)
PDEmut
EndoUmut

Figure S12

**Table 1A. Dinucleotide enrichment and de-enrichment in WT BMM at 12 hpi(position -2 to -1)**

| | MHV(S) | | MHV(V) | | PDEmut | | EndoUmut | |
|---|---|---|---|---|---|---|---|---|
| Dinucleotide | Fold change | qval | Fold change | qval | Fold change | qval | Fold change | qval |
| UA | -0.57 | 1 | -0.59 | 1 | 1.36 | 0.0E-1000 | 1.71 | 0.0E-1000 |
| UU | 1.01 | 0.0E-1000 | 0.97 | 0.0E-1000 | 1.22 | 0.0E-1000 | 1.35 | 0.0E-1000 |
| UG | -1.87 | 1 | -1.98 | 1 | -0.69 | 1 | -0.29 | 1 |
| UC | 1.80 | 0.0E-1000 | 1.99 | 0.0E-1000 | 0.87 | 4.81E-123 | 0.15 | 0.0018 |
| GA | -1.65 | 1 | -1.79 | 1 | -1.24 | 1 | -1.05 | 1 |
| GU | 0.29 | 1.19E-23 | 0.15 | 6.35E-07 | -0.23 | 1 | -0.94 | 1 |
| GG | -2.67 | 1 | -2.83 | 1 | -2.65 | 1 | -2.70 | 1 |
| GC | 0.69 | 5.32E-99 | 0.78 | 1.22E-120 | -0.30 | 1 | -1.01 | 1 |
| CA | -0.89 | 1 | -1.00 | 1 | -0.42 | 1 | 0.20 | 6.52E-07 |
| CU | -0.11 | 1 | -0.27 | 1 | -0.55 | 1 | -0.88 | 1 |
| CG | -1.94 | 1 | -2.05 | 1 | -1.98 | 1 | -1.86 | 1 |
| CC | 0.52 | 3.96E-38 | 0.62 | 4.42E-52 | -0.45 | 1 | -1.03 | 1 |
| AA | -1.60 | 1 | -1.77 | 1 | -1.01 | 1 | -0.74 | 1 |
| AU | 0.10 | 0.0005 | 0.00 | 1 | -0.34 | 1 | -0.90 | 1 |
| AG | -2.20 | 1 | -2.44 | 1 | -2.11 | 1 | -2.08 | 1 |
| AC | 0.98 | 3.21E-174 | 1.11 | 7.50E-221 | 0.10 | 0.0195 | -0.61 | 1 |

**Table 1B. Dinucleotide enrichment and de-enrichment in WT BMM at 9 hpi(position -2 to -1)**

| | MHV(S) | | MHV(V) | | PDEmut | | EndoUmut | |
|---|---|---|---|---|---|---|---|---|
| Dinucleotide | Fold change | qval | Fold change | qval | Fold change | qval | Fold change | qval |
| UA | -0.20 | 1 | 0.13 | 0.00135 | 0.19 | 0.0071 | 1.43 | 0.0E-1000 |
| UU | 1.04 | 0.0E-1000 | 1.21 | 0.0E-1000 | 0.94 | 9.92E-88 | 1.27 | 0.0E-1000 |
| UG | -1.71 | 1 | -1.43 | 1 | -1.21 | 1 | -0.45 | 1 |
| UC | 1.57 | 0.0E-1000 | 0.92 | 4.41E-84 | 0.70 | 8.77E-17 | 0.68 | 5.39E-59 |
| GA | -1.22 | 1 | -0.94 | 1 | -0.88 | 1 | -1.03 | 1 |
| GU | 0.41 | 4.91E-41 | 0.50 | 8.87E-43 | 0.65 | 2.73E-27 | -0.60 | 1 |
| GG | -2.46 | 1 | -2.11 | 1 | -1.51 | 1 | -2.78 | 1 |
| GC | 0.50 | 1.95E-41 | 0.01 | 0.981 | 0.26 | 0.00181 | -0.41 | 1 |
| CA | -0.71 | 1 | -0.53 | 1 | -0.15 | 1 | 0.21 | 2.18E-07 |
| CU | -0.03 | 1 | 0.08 | 0.111 | 0.29 | 0.00019 | -0.81 | 1 |
| CG | -1.72 | 1 | -1.40 | 1 | -1.27 | 1 | -1.93 | 1 |
| CC | 0.29 | 1.19E-10 | -0.11 | 1 | -0.09 | 1 | -0.49 | 1 |
| AA | -1.22 | 1 | -0.89 | 1 | -0.98 | 1 | -0.77 | 1 |
| AU | 0.12 | 0.00019 | 0.31 | 2.33E-16 | 0.32 | 6.74E-07 | -0.79 | 1 |
| AG | -1.95 | 1 | -1.76 | 1 | -1.97 | 1 | -2.28 | 1 |
| AC | 0.66 | 6.51E-63 | 0.17 | 0.0014 | -0.03 | 1 | -0.12 | 1 |

Table S1

#### Table 2A. Dinucleotide enrichment and de-enrichment in WT BMM at 12 hpi(position -1 to +1)

| | MHV$^{(S)}$ | | MHV$^{(V)}$ | | PDE$^{mut}$ | | EndoU$^{mut}$ | |
|---|---|---|---|---|---|---|---|---|
| **Dinucleotide** | **Fold change** | **qval** | **Fold change** | **qval** | **Fold change** | **qval** | **Fold change** | **qval** |
| UA | 2.00 | 0.0E-1000 | 1.93 | 0.0E-1000 | 1.43 | 0.0E-1000 | 1.02 | 1.05E-281 |
| UU | -0.85 | 1 | -1.06 | 1 | -0.19 | 1 | -0.18 | 1 |
| UG | -0.82 | 1 | -0.79 | 1 | -0.06 | 1 | 0.15 | 3.42E-07 |
| UC | -0.67 | 1 | -0.94 | 1 | -0.99 | 1 | -0.84 | 1 |
| GA | -1.94 | 1 | -2.03 | 1 | -1.17 | 1 | -0.93 | 1 |
| GU | -1.81 | 1 | -1.97 | 1 | -1.04 | 1 | -0.84 | 1 |
| GG | -2.60 | 1 | -2.75 | 1 | -1.44 | 1 | -0.85 | 1 |
| GC | -2.39 | 1 | -2.59 | 1 | -2.50 | 1 | -2.34 | 1 |
| CA | 2.46 | 0.0E-1000 | 2.60 | 0.0E-1000 | 1.40 | 0.0E-1000 | 0.56 | 8.72E-50 |
| CU | -1.01 | 1 | -0.95 | 1 | -1.31 | 1 | -1.57 | 1 |
| CG | -0.58 | 1 | -0.28 | 1 | -1.00 | 1 | -1.38 | 1 |
| CC | -0.49 | 1 | -0.35 | 1 | -1.32 | 1 | -1.82 | 1 |
| AA | -0.83 | 1 | -0.89 | 1 | 0.51 | 1.78E-68 | 0.98 | 2.08E-239 |
| AU | -0.88 | 1 | -1.00 | 1 | 0.37 | 3.80E-39 | 0.62 | 5.98E-95 |
| AG | -1.36 | 1 | -1.44 | 1 | 0.04 | 0.670003 | 0.41 | 8.99E-31 |
| AC | -1.64 | 1 | -1.73 | 1 | -0.72 | 1 | -0.37 | 1 |

#### Table 2B. Dinucleotide enrichment and de-enrichment in WT BMM at 9 hpi(position -1 to -1)

| | MHV$^{(S)}$ | | MHV$^{(V)}$ | | PDE$^{mut}$ | | EndoU$^{mut}$ | |
|---|---|---|---|---|---|---|---|---|
| **Dinucleotide** | **Fold change** | **qval** | **Fold change** | **qval** | **Fold change** | **qval** | **Fold change** | **qval** |
| UA | 2.01 | 0.0E-1000 | 2.06 | 0.0E-1000 | 2.08 | 0.0E-1000 | 1.28 | 0.0E-1000 |
| UU | -0.91 | 1 | -0.52 | 1 | -0.76 | 1 | -0.46 | 1 |
| UG | -0.41 | 1 | -0.04 | 1 | -0.28 | 1 | -0.02 | 1 |
| UC | -0.72 | 1 | -0.79 | 1 | -0.59 | 1 | -0.84 | 1 |
| GA | -1.61 | 1 | -1.27 | 1 | -1.59 | 1 | -1.05 | 1 |
| GU | -1.70 | 1 | -1.48 | 1 | -1.24 | 1 | -1.10 | 1 |
| GG | -2.24 | 1 | -2.00 | 1 | -1.73 | 1 | -1.00 | 1 |
| GC | -2.36 | 1 | -2.03 | 1 | -1.38 | 1 | -2.23 | 1 |
| CA | 2.18 | 0.0E-1000 | 1.49 | 0.0E-1000 | 1.38 | 3.23E-112 | 1.24 | 2.27E-302 |
| CU | -1.11 | 1 | -1.17 | 1 | -0.97 | 1 | -1.61 | 1 |
| CG | -0.41 | 1 | -0.35 | 1 | -0.42 | 1 | -0.96 | 1 |
| CC | -0.70 | 1 | -0.98 | 1 | -0.90 | 1 | -1.40 | 1 |
| AA | -0.35 | 1 | -0.06 | 1 | 0.03 | 1 | 0.86 | 3.03E-182 |
| AU | -0.69 | 1 | -0.40 | 1 | -0.42 | 1 | 0.32 | 2.80E-24 |
| AG | -0.99 | 1 | -0.66 | 1 | -0.59 | 1 | 0.27 | 1.19E-13 |
| AC | -1.40 | 1 | -1.15 | 1 | -0.65 | 1 | -0.45 | 1 |

Table S2

| Position (bp) | Reference | Variant |
|---|---|---|
| 67 | T | C |
| 256 | A | T |
| 2471 | T | C |
| 5304 | T | C |
| 6265 | C | T |
| 6796 | A | T |
| 7475 | G | A |
| 9245 | C | T |
| 10136 | A | C |
| 12140 | G | A |
| 13604 | G | A |
| 16624 | G | C |
| 16990 | C | T |
| 17533 | T | G |
| **18646** | **A** | **C** |
| 18681 | C | T |
| **20552** | **C** | **G** |
| **20553** | **A** | **C** |
| **20554** | **T** | **G** |
| **22147** | **A** | **G** |
| 22739 | T | G |
| 22742 | T | C |
| 23393 | C | T |
| 23860 | T | C |
| 24404 | T | A |
| 24447 | G | A |
| 24448 | C | T |
| 24450 | T | A |
| 26074 | G | C |
| **28960** | **T** | **C** |
| 29012 | G | C |
| 29650 | T | C |

Table S3
